# Systematic Survey of Public Datasets for Behavioral Research in Invertebrate Models: Toward FAIR and Standardized Data Sharing

**DOI:** 10.64898/2025.12.12.693879

**Authors:** Natalia Piórkowska, Róża Mazurek, Damian Adamek, Martyna Łopianiak, Mikołaj Kubś, Krzysztof Kulka, Patryk Łuszczek, Radosław Mijał, Alan Ostromęcki

## Abstract

Behavioral datasets for invertebrate model organisms are rapidly expanding alongside automated imaging, tracking, and artificial intelligence (AI) based phenotyping, yet their technical structure and compliance with the Findable, Accessible, Interoperable and Reusable (FAIR) principles remain heterogeneous. We present a two-stage survey of openly available behavioural datasets for major invertebrate models *Caenorhabditis elegans (C. elegans)*, *Drosophila melanogaster (D. melanogaster)*, *Galleria mellonella* (*G. mellonella*), and planarians *Schmidtea mediterranea* (*S. mediterranea*) with larval zebrafish (*Danio rerio)* included as a vertebrate comparator. Stage 1 comprised a PRISMA-guided literature review (from 2015 to 2025) across indexed databases and complementary non-indexed sources, yielding 12 eligible publications describing 12 open behavioural datasets. Stage 2 independently screened and technically evaluated repository deposits (from June 2022 to July 2025), producing a final corpus of 20 datasets scored on a four-dimension ordinal rubric capturing usability, annotation richness, technical quality and AI-readiness. All extracted descriptors, repository search logs, and scoring sheets are released as public data records enabling full regeneration of figures and summary statistics. Across Stage 2 deposits, multimodality and open file formats were common, whereas interoperability and AI-readiness were most constrained by limited machine-readable metadata, weak raw-to-derived provenance, and sparse adoption of formal standards or ontologies. This Data Descriptor provides a reproducible, dataset-centred overview of behavioural resources for invertebrate models and practical guidance for FAIR-aligned publication, secondary biological analyses, and AI benchmarking.

## Background & Summary

Behavioural research in invertebrate model organisms—including *C. elegans*, *D. melanogaster*, *G. mellonella*, and planarians such as *S. mediterranea*—supports high-throughput dissection of genetic, neural, and ecological determinants of behaviour.[1–4] Advances in automated imaging, posture tracking, and machine-learning–assisted behavioural phenotyping have accelerated the generation of large-scale behavioural resources, including high-throughput tracking platforms and deep-learning–based pose estimation.[1–2, 5–7, 12] These datasets increasingly comprise multimodal deposits that combine raw video or imaging with derived trajectories, pose outputs, time-resolved feature matrices, and behavioural annotations.[5, 7, 9]

The shift toward open behavioural resources is tightly linked to the FAIR (Findable, Accessible, Interoperable, Reusable) principles.[8] FAIR-aligned sharing requires persistent identifiers, transparent licensing, rich human- and machine-readable metadata, and interoperable file formats, supported by community standards and data languages that explicitly target machine actionability.[8–10] Behavioural datasets are often complex and highly dependent on experimental context; without consistent metadata structures and clear provenance between raw and derived layers, reanalysis, cross-laboratory comparison, and reuse for AI benchmarking remain difficult. Existing invertebrate datasets vary widely in file organisation, metadata depth, annotation schemas, licensing, and repository practices. Machine-readable codebooks, formal standards, and ontology mappings remain intermittent, limiting computational interoperability despite emerging FAIR-focused infrastructures in neuroscience.[9–10]

To our knowledge, no technically oriented survey has systematically mapped publicly available behavioural datasets for invertebrate models in terms of data structures, metadata practices, repository dissemination, FAIR implementation, and standards adoption. Here we provide a reproducible two-stage survey of open behavioural datasets for major invertebrate models, with larval zebrafish *Danio rerio* included as a structured comparator reflecting a vertebrate ecosystem with mature curated resources.

Across both stages we address five interrelated research questions:

**RQ1:** What data modalities, file formats, and organisational structures are used, and what are their implications for interoperability?

**RQ2:** How complete and structured are metadata describing organism identity, experimental conditions, and acquisition/processing pipelines?

**RQ3:** How are datasets disseminated (repositories, licensing, documentation, APIs or scripts), and how does this affect reuse?

**RQ4:** To what extent do datasets implement FAIR dimensions through persistent identifiers, open formats, structured metadata, and licensing?

**RQ5:** What is the level of adoption of formal standards and ontologies, and what emerging harmonisation efforts are evident?

By integrating PRISMA-based bibliographic evidence with repository-level technical screening,[11] this Data Descriptor delivers a dataset-centred overview of behavioural resources for invertebrate models and provides practical recommendations for FAIR-aligned behavioural data publication.

## Methods

### Study overview

This study comprised two complementary but methodologically independent components addressing publicly available behavioural datasets. The primary focus was on model invertebrates (*D. melanogaster*, *C. elegans*, *G. mellonella*, and *S. mediterranea*), with larval zebrafish (*Danio rerio*) included to facilitate cross-taxon comparison of data-sharing and metadata practices. Stage 1 was a PRISMA 2020–compliant systematic literature review designed to identify dataset descriptor publications. Stage 2 was an independent technical evaluation of datasets through direct repository inspection. No new experimental data were generated; all analyses were based on publicly available datasets and associated metadata.

### Stage 1: Systematic literature review (PRISMA 2020)

#### Information sources and search

We searched PubMed/MEDLINE, Scopus, Web of Science Core Collection, IEEE Xplore, and the ACM Digital Library for English-language records published between 1 January 2015 and 30 June 2025. Indexed database searches were complemented by screening (i) the first 200 Google Scholar results, (ii) relevant bioRxiv and arXiv preprints, and (iii) reference lists of all included papers. Full database-specific strategies, interfaces/providers, run dates, time windows, platform-specific filters, and retrieved counts are reported in Supplementary Table S1 in accordance with PRISMA-S guidance.

#### Eligibility criteria

Records were eligible if they (i) described publicly accessible behavioural datasets from target invertebrate models or larval zebrafish *Danio rerio*, (ii) reported technical information on data structure or documentation (file formats, folder organisation, metadata, annotations), and (iii) provided verifiable dataset access (DOI, accession, or persistent link). During title/abstract screening, records were assessed for taxon relevance, language, and time window, whereas technical detail requirements were applied at the full-text stage (Table 1). The review protocol was not prospectively registered.

**Table 1.** Eligibility criteria for Stage 1 systematic literature review (PRISMA 2020).

| Screening level | Inclusion criteria | Exclusion criteria |
| --- | --- | --- |
| **Title/abstract screening** | (i) Peer-reviewed articles, conference proceedings, or preprints describing a publicly available behavioral dataset for target invertebrate models (*D. melanogaster*, *C. elegans*, *G. mellonella*, *S. mediterranea*) or larval *Danio rerio* (technical details not required at abstract stage); (ii) published in English; (iii) published between 2015-01-01 and 2025-06-30. | i) Vertebrate-only datasets (except larval *Danio rerio*); (ii) non-model species only / unrelated systems; (iii) behavioral studies without an open dataset; (iv) non-scholarly sources (blogs, popular science, informal notes); (v) non-English records or outside the time window. |
| **Full-text eligibility** | (i) Full text accessible; (ii) provides technical dataset information, e.g., data structure/format, metadata schema/content, annotations, repository/DOI, license, FAIR or standardization statements; (iii) dataset link/DOI/accession verifiable. | (i) Full text unavailable; (ii) insufficient technical detail on the dataset; (iii) purely theoretical/conceptual papers without dataset description or evaluation; (iv) technical/IT reports not tied to an empirical behavioral dataset; (v) methods/ontology papers without linkage to a specific dataset or case study. |

#### Screening and selection

All records were imported into Mendeley Reference Manager (v2.115). After automated and manual de-duplication, two reviewers (RM, AO) independently screened titles/abstracts and full texts using pre-specified criteria (Table 1). Disagreements were resolved by consensus, with a third reviewer (NP) arbitrating unresolved cases.

A PRISMA 2020 flow diagram is provided as Figure 1, and source-resolved inclusion and exclusion counts are summarised in Supplementary Table S11.

**Figure 1.**
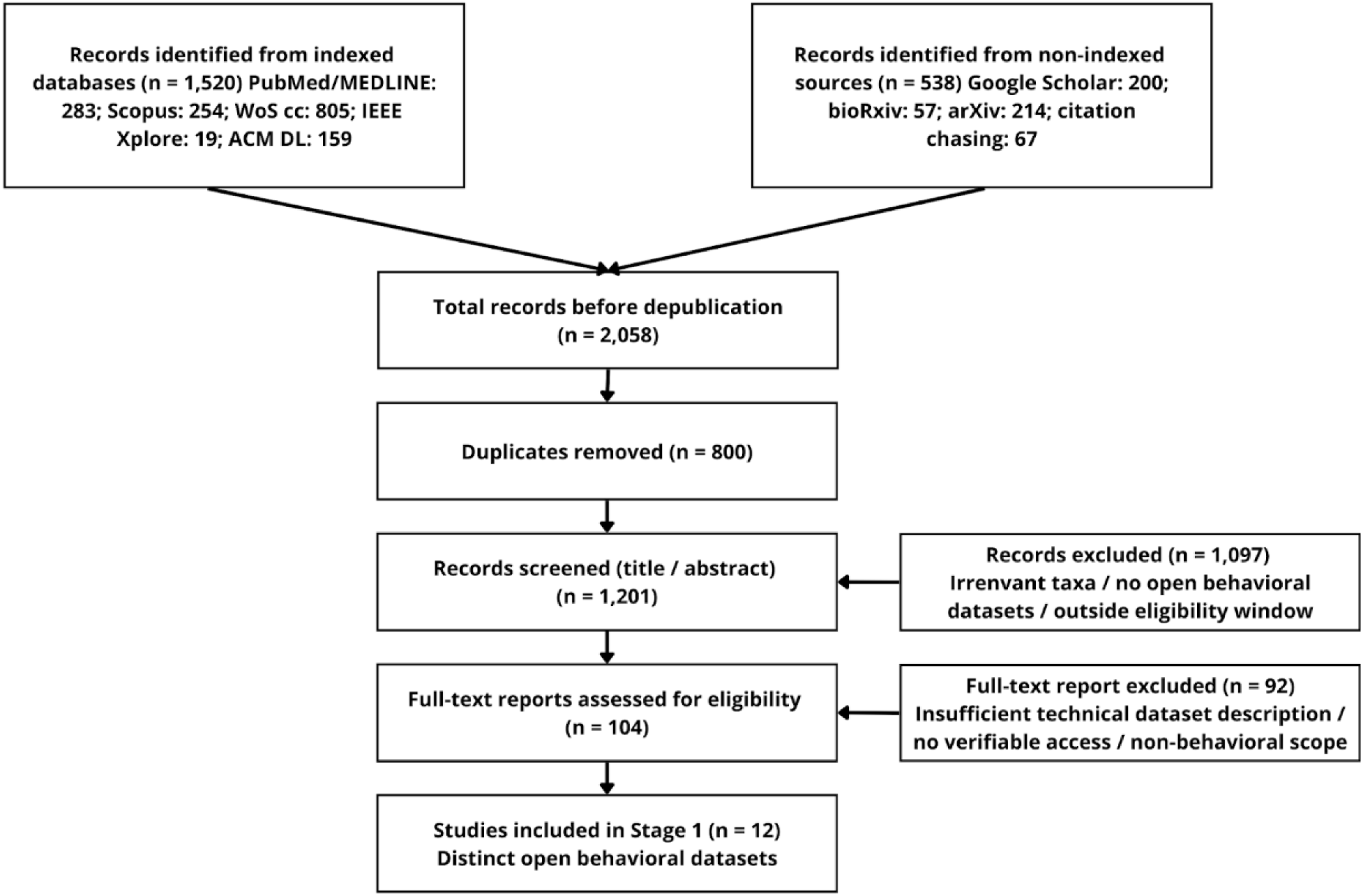
PRISMA 2020 flow diagram.

#### Data extraction and synthesis

A piloted extraction template captured bibliographic variables and dataset-level descriptors aligned with RQ1–RQ5. Stage 1 extraction recorded descriptors explicitly reported in publications, whereas Stage 2 extraction verified and expanded these descriptors through direct inspection of repository deposits. Missing or ambiguous information was coded explicitly as “Not reported”, and non-applicable items as “NA”.

#### Quality and bias assessment

Reporting/accessibility bias was evaluated at record level using a three-dimension rubric scoring documentation completeness, metadata/FAIR/licensing transparency, and standards/ontologies reporting on 0–2 points scales (Supplementary Table S5a). Dimension scores were summed (0–6) and mapped to High/Moderate/Low record-quality categories (Supplementary Table S5) using predefined thresholds (Supplementary Table S5b). Two reviewers rated all records independently.

### Stage 2: Technical screening and evaluation of datasets

#### Repository search and screening

Major general-purpose repositories (Zenodo, Figshare, Dryad, OSF, IEEE DataPort, Harvard Dataverse) and specialised archives were considered. Searches were executed iteratively between 4 July 2025 and 29 July 2025, targeting deposits from June 2022–July 2025. Queries were built from predefined organism-, modality/behaviour-, FAIR/accessibility-, and dataset-type keyword families (Supplementary Table S6) and adapted to each repository’s syntax. Where archives lacked functional dataset search interfaces at the time of screening, they were recorded but not systematically queried. Records were screened for behavioural relevance, open accessibility, minimum dataset-level metadata, and suitability for technical evaluation. A summarised repository screening log is provided in Supplementary Table S7; the full audit trail of executed searches and retrieved counts is available as Supplementary Data File S2 (CSV).

#### Dataset evaluation protocol

Eligible datasets were downloaded between 15 and 30 July 2025 using the most recent stable DOI-linked version; for deposits without DOI, the latest tagged release/commit was used. Files were inspected for internal consistency and scored with the technical extraction rubric (Supplementary Table S8). Dataset descriptors were extracted using a pre-specified variable dictionary (Supplementary Table S10) covering identification, taxa, modality/structure, metadata, sharing/FAIR indicators, and AI-readiness. Four technical dimensions were scored on 1–3 ordinal scales: Usability, Annotation richness, Technical quality, and AI-readiness (Supplementary Table S8). Two reviewers (AO, NP) scored all datasets independently using masked identifiers. Inter-rater agreement was quantified per dataset using Cohen’s κ; κ categories followed Landis–Koch thresholds. Per-dataset R1/R2 scores and κ estimates are reported in Supplementary Table S9.

#### Synthesis and visualisation

Categorical variables were summarised as frequencies; ordinal scores were analysed as ranked distributions. Visualisations were generated in Python 3.10 (pandas, matplotlib).

### Data Records

All data products generated in this survey are released as Supplementary Information to enable independent verification, re-analysis, and reuse. The released materials cover both stages—Systematic literature review (Stage 1) and repository-based dataset discovery with technical scoring (Stage 2)—as well as all dataset-level descriptors and derived summaries.

The complete Stage 2 master extraction sheet (Supplementary Data File S1, CSV/XLSX) contains dataset-level descriptors for DS01–DS20 and serves as the direct source for Figures 3–6. Binary modality and file-format matrices, FAIR checklists, and rater-level scoring sheets with κ estimates are provided as dedicated tabs within the same file. Stage 1 outputs are provided as three complementary extraction tables (Supplementary Tables S2–S4) aligned with RQ1–RQ5, and Stage 1 search strategies and bias scoring rubrics are reported in Supplementary Tables S1 and S5a–S5b. The full Stage 2 repository audit trail is released as Supplementary Data File S2.

## Results

### 3.1 Results of Stage 1 (PRISMA literature review)

#### PRISMA flow summary

Database searches retrieved n = 1,520 records across indexed sources (PubMed/MEDLINE, Scopus, Web of Science Core Collection, IEEE Xplore, ACM Digital Library). Screening of non-indexed sources added n = 200 Google Scholar hits, n = 57 bioRxiv preprints, n = 214 arXiv preprints, and n = 67 records from reference list screening, yielding n = 2,058 records in total. After de-duplication, n = 800 duplicates were removed, leaving n = 1,201 unique records for title/abstract screening. At this stage, n = 1,097 records were excluded as irrelevant to target taxa, lacking openly accessible behavioural datasets, or falling outside the eligibility window. Full texts of n = 104 records were assessed; n = 92 were excluded due to insufficient technical dataset description, lack of verifiable access, or non-behavioural scope. The final Stage 1 corpus comprised n = 12 eligible publications, each corresponding to a distinct openly accessible behavioural dataset (Figure 1).

#### Characteristics of included records

Eligible publications spanned 2015–2025, with a strong concentration in later years reflecting the rapid growth of automated phenotyping, high-throughput tracking, and repository-based sharing in behavioural research. The overall annual counts of included records show a clear increase in the second half of the observation window (Figure 2), consistent with broader trends in open data and AI-assisted behavioural analysis.

**Figure 2.**
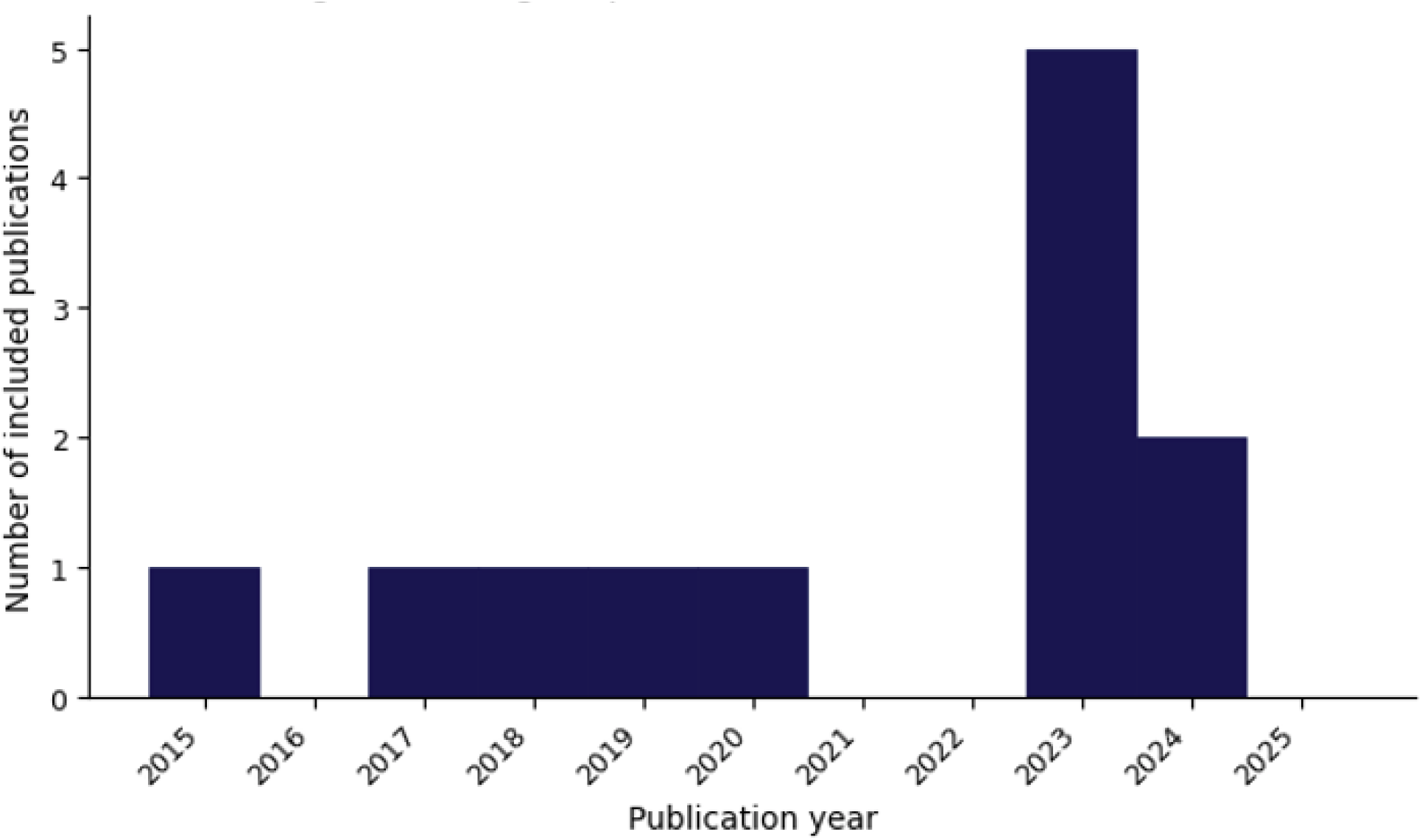
Stage 1 temporal distribution of included records (2015–2025).

Beyond the aggregate trend, Stage 1 records were unevenly distributed across taxa. Figure 3 presents the same temporal distribution stratified by model organism, highlighting how contributions from *C. elegans*, *D. melanogaster*, *G. mellonella*, planarians, and larval *Danio rerio* are staggered over time. This taxon-resolved view reveals differences in the timing and intensity of dataset publication between vertebrate and invertebrate ecosystems and emphasises that technically detailed, openly shared behavioural datasets remain concentrated in a relatively small number of model-specific “pockets”.

**Figure 3.**
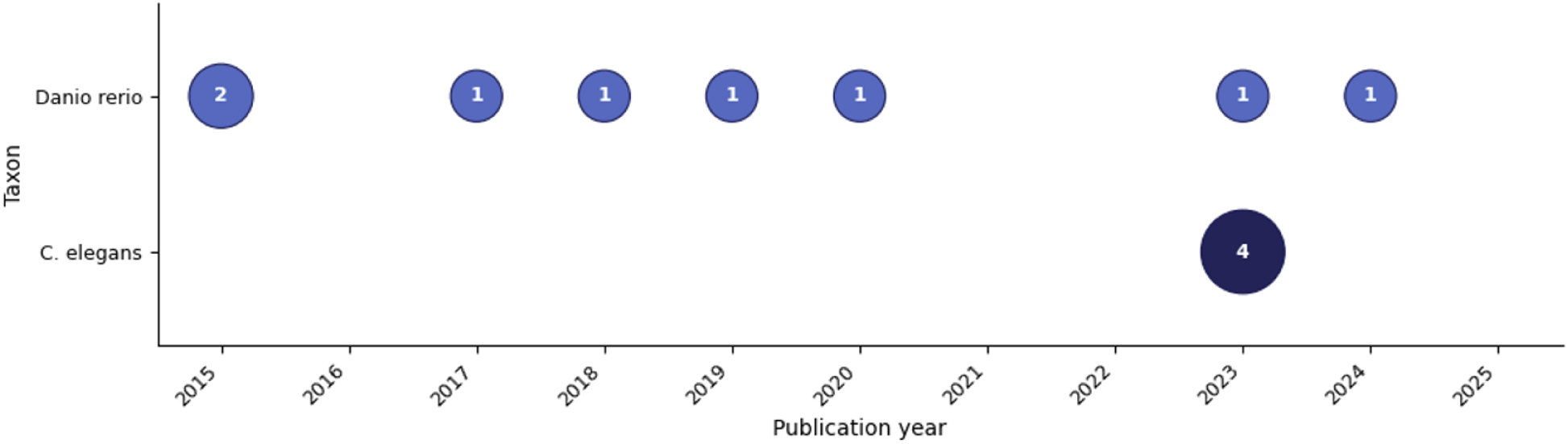
Stage 1 temporal distribution of included records by taxon (2015–2025).

#### Stage 1 evidence mapped to RQ1–RQ5

##### RQ1 (data structures and formats)

Most Stage 1 records described multimodal datasets coupling raw imaging/video with derived trajectories or pose estimates and/or feature matrices. Open, analysis-ready formats predominated (e.g. CSV/TSV, standard video containers), whereas explicit file-level provenance between raw and derived layers was inconsistently reported. In several cases, raw and processed files were co-located in shared directories without clear linking conventions, complicating reproducible reanalysis.

##### RQ2 (metadata reporting)

Metadata completeness varied widely. Human-readable documentation (method sections, README files) was common, but structured machine-readable schemas, variable dictionaries, and formal metadata standards were rare. Typical gaps included partial organism descriptors, incomplete reporting of experimental conditions, and minimal information on processing pipelines, limiting computational interpretability and cross-dataset comparability.

##### RQ3 (dissemination practices)

Dissemination relied primarily on general-purpose repositories supporting stable landing pages and DOI assignment, complemented by hybrid hosting (repository + GitHub) and, less frequently, GitHub-only deposits. While most records linked to at least one persistent identifier, dataset-level licensing information was sometimes missing or embedded only in article text, creating ambiguity about permissible reuse.

##### RQ4 (FAIR framing)

FAIR implementation was skewed toward Findability and Accessibility, supported through repository indexing, stable DOIs, and human-readable landing pages. Interoperability and Reusability were least often operationalised: structured metadata, machine-readable codebooks, and ontology alignment were rarely present. Overall, most Stage 1 datasets were FAIR-Partial; FAIR-Comprehensive support was largely confined to curated *Danio rerio* resources.

##### RQ5 (standards/ontologies)

Formal standards and ontologies were rarely referenced in invertebrate datasets. Custom label sets without explicit mapping to shared vocabularies predominated, while zebrafish resources more frequently leveraged existing ontology stacks and standard identifiers. This asymmetry underscores the lack of harmonised behavioural data standards in invertebrate ecosystems.

#### Reporting/accessibility bias

Record-level reporting quality was heterogeneous. Moderate and low ratings were driven mainly by incomplete technical descriptions of data structures, weak raw-to-derived traceability, unclear licensing, and limited machine-readable metadata. Using the Stage 1 bias-score thresholds (0–6), records stratified into High (5–6), Moderate (3–4), and Low (0–2) categories (Figure 4). This heterogeneity indicates that even datasets positioned as open resources frequently fall short of reproducible, machine-actionable

**Figure 4.**
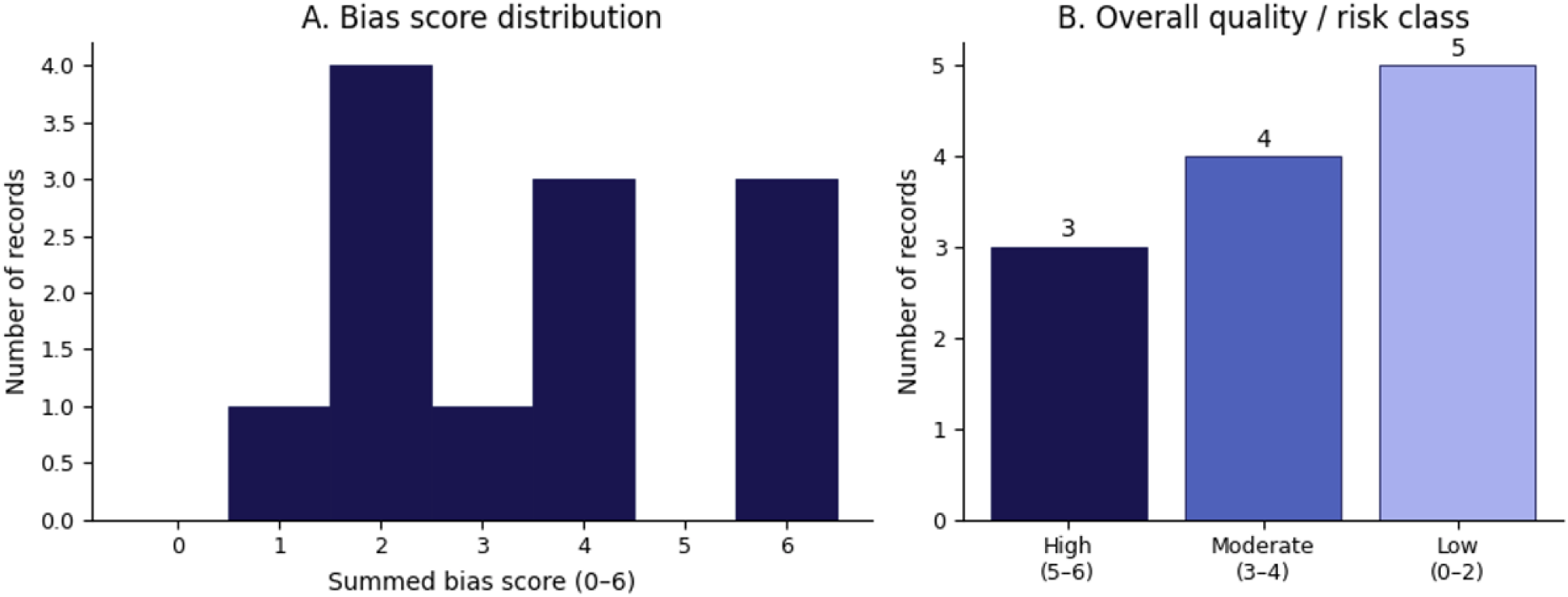
Distribution of Stage 1 reporting/accessibility bias scores.

#### Dissemination landscape by taxon and FAIR class

Mapping taxon → sharing platform → FAIR class showed clear ecosystem-level differences. *C. elegans* deposits flowed primarily into GitHub-only and other non-curated platforms and culminated predominantly in FAIR-Partial classification. In contrast, *Danio rerio* records showed a strong stream into ZFIN and related curated infrastructures, consistently mapping to FAIR-Comprehensive. Hybrid hosting (e.g. Zenodo + GitHub) was distributed across FAIR classes, indicating that DOI assignment and repository deposition improve Findability and Accessibility but do not guarantee interoperability or rich reuse (Figure 5A,B). These patterns highlight a structural gap between mature, ontology-backed vertebrate infrastructures and more fragmented invertebrate sharing practices.

**Figure 5.**
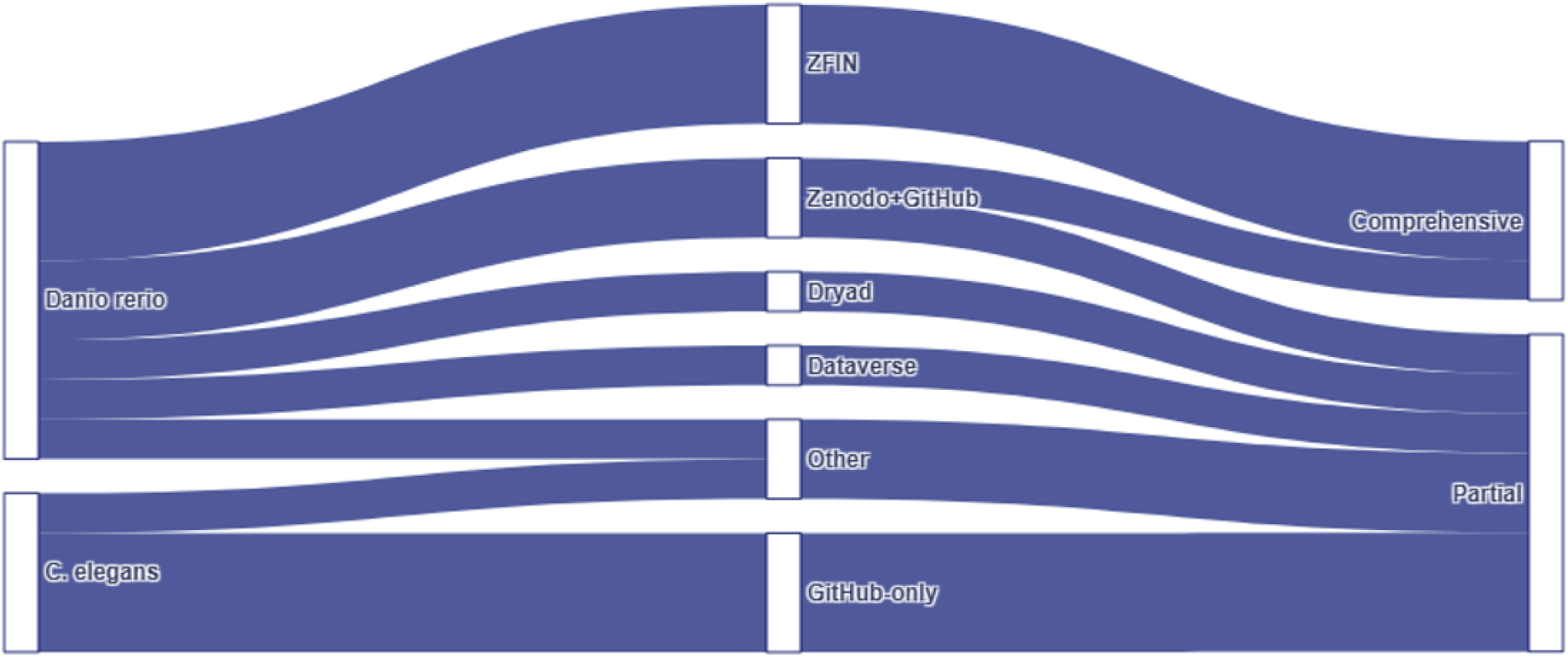
A. Stage 1 dissemination landscape: Taxon → Platform → FAIR class.

**Figure 5B.**
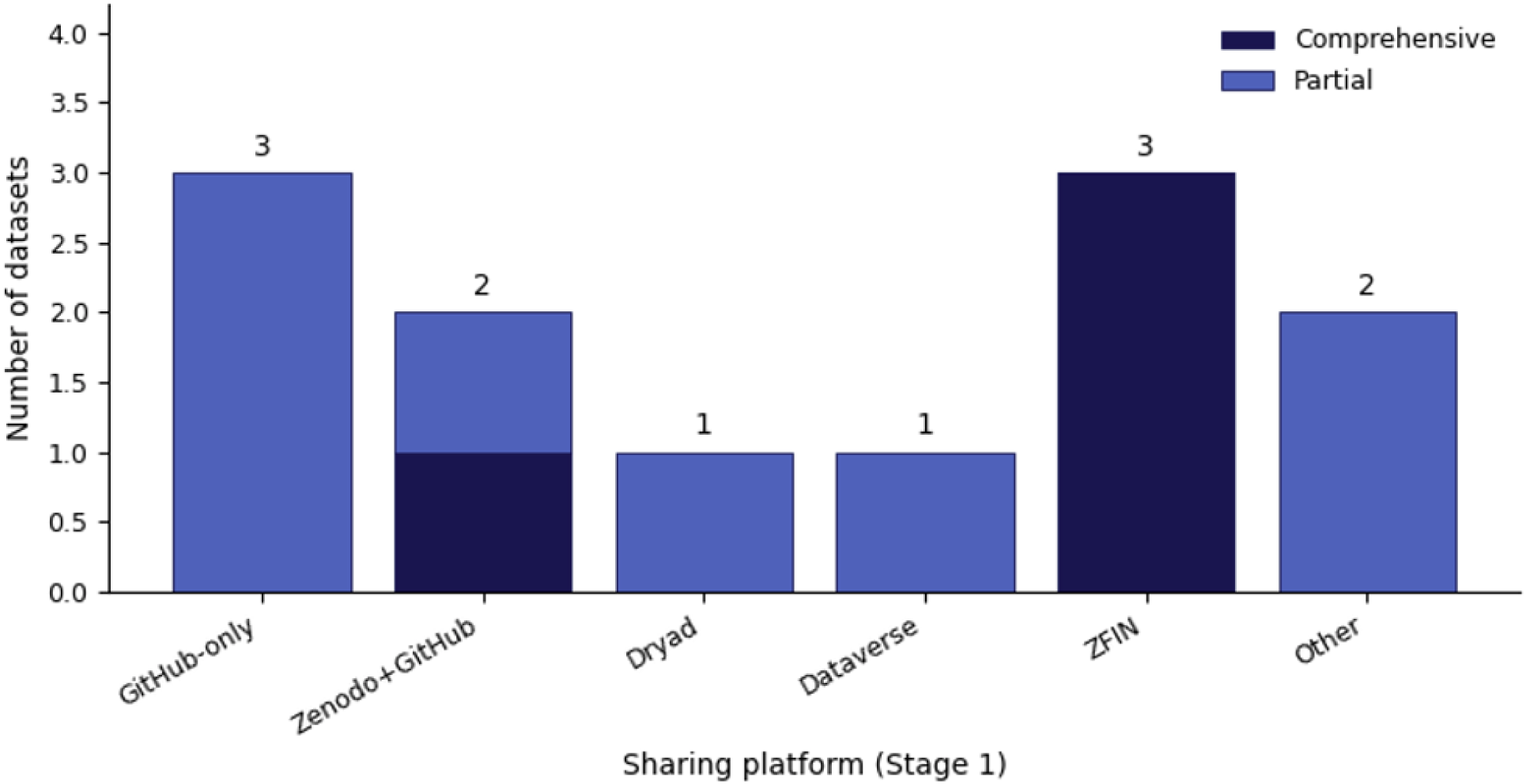
Stage 1 repository & dissemination landscape by FAIR class.

### 3.2 Results of Stage 2 (technical screening)

#### Final Stage 2 dataset set

Stage 2 screening identified n = 20 eligible datasets (DS01–DS20) spanning target invertebrate models and a larval zebrafish comparator. Deposits lacking verifiable access, behavioural relevance, or minimum dataset-level metadata were excluded prior to scoring. A condensed overview of included datasets, including taxa, modalities, access routes, and licensing, is provided in Table 2.

**Table 2.** Stage 2 dataset overview.

| Dataset | Organism | Repository/platform | Modalities | DOI/PID | License clarity | Mean technical score |
| --- | --- | --- | --- | --- | --- | --- |
| WormSwin: *C. elegans* Video Datasets (DS01) | *C. elegans* | Zenodo | Video, ImagesFrames | yes | yes | 2.38 |
| Three-dimensional body reconstruction enables quantification of liquid consumption in small invertebrates (DS02) | *Linepithema humile* | Zenodo | Video, ImagesFrames, TrackingTrajectories, PoseSkeleton, FeatureMatrices | yes | yes | 2.25 |
| Concatenated planarian behavioral barcodes (DS03) | *Dugesia japonica* | Figshare | FeatureMatrices | yes | yes | 2.00 |
| Differential neuroanatomical, neurochemical, and behavioral impacts of early-age isolation in a eusocial insect (DS04) | *Camponotus floridanus* | Dryad | FeatureMatrices | yes | yes | 2.00 |
| Mantis shrimp locomotion: coordination and variation of hybrid metachronal swimming (DS05) | *Neogonodactylus bredini* | Dryad | Video, FeatureMatrices | yes | yes | 2.88 |
| Multifaceted and extensive behavioral trajectories of genomically diverse *Drosophila* lines (DS06) | *Drosophila melanogaster* | Dryad | Video, TrackingTrajectories, FeatureMatrices | yes | yes | 2.38 |
| Miniature linear and split-belt treadmills reveal mechanisms of adaptive motor control in walking *Drosophila* ((DS07)) | *Drosophila melanogaster* | Dryad | TrackingTrajectories, FeatureMatrices | yes | yes | 2.38 |
| Asynchronous haltere input drives specific wing and head movements in *Drosophila* (DS08) | *Drosophila melanogaster* | Dryad | Video, TrackingTrajectories, FeatureMatrices | yes | yes | 2.38 |
| 2D Tracking and 3D reconstruction of legs from tethered walking *Drosophila* (DS09) | *Drosophila melanogaster* | Dataverse | Video, TrackingTrajectories, FeatureMatrices | yes | yes | 2.12 |
| Crawling Trajectories of *Drosophila* Larvae Responding to Vibration (DS10) | *Drosophila melanogaster* (larvae) | Dataverse | Video, TrackingTrajectories, FeatureMatrices | yes | yes | 2.38 |
| Individual flight headings for *Drosophila melanogaster* in response to different wavelengths of light (DS11) | *Drosophila melanogaster* | Dryad | FeatureMatrices | yes | yes | 1.50 |
| Neural mechanisms to incorporate visual counterevidence in self movement estimation (DS12) | *Drosophila melanogaster* | Dryad | FeatureMatrices | yes | yes | 1.75 |

**Table 2.** Stage 2 dataset overview.
|  |  |  |  |  |  |  |
| --- | --- | --- | --- | --- | --- | --- |
| Zebrafish Seizure Behavior Classification Dataset and Trained ML Models (DS13) | Danio rerio (larvae) | Zenodo | Video, TrackingTrajectories, FeatureMatrices | yes | yes | 2.50 |
| Chloride-dependent mechanisms of multimodal sensory discrimination and nociceptive sensitization in Drosophila (DS14) | Drosophila melanogaster | Dryad | FeatureMatrices | yes | yes | 2.38 |
| Dimensionality of locomotor behaviors in developing C. elegans (DS15) | C. elegans | Dryad | TrackingTrajectories, FeatureMatrices | yes | yes | 2.00 |
| An improved neural network model enables worm tracking in challenging conditions and increases SNR in phenotypic screens (datasets and models) (DS16) | C. elegans | Zenodo | TrackingTrajectories | yes | yes | 2.62 |
| Locomotion of naive Drosophila larvae depending on age (DS17) | Drosophila melanogaster (larvae) | GIN / G-Node | Video, TrackingTrajectories | yes | yes | 1.50 |
| Zebrafish larvae exploration and aversive chemotaxis dataset (DS18) | Danio rerio (larvae) | Dryad | TrackingTrajectories | yes | yes | 2.38 |
| Sensory neurons couple arousal and foraging decisions in C. elegans (DS19) | C. elegans | Dryad | TrackingTrajectories | yes | yes | 2.25 |
| Eigenworms (DS20) | C. elegans | Project-specific | TimeSeries, TrackingTrajectories | yes | no | 2.12 |

#### Stage 2 evidence mapped to RQ1–RQ5

##### RQ1 (modalities, formats, organisation)

Stage 2 deposits were predominantly multimodal, combining raw recordings with derived quantitative layers. Tracking trajectories were the most frequent modality (13/20 datasets), closely followed by tabular or feature matrices (12/20), while raw video was present in 9/20 deposits (Figure 6). Image-frame archives and neural or other signal layers were rare (2/20 each), and explicit pose-skeleton outputs appeared once (1/20). This distribution suggests that, in practice, many invertebrate datasets are shared primarily as trajectory and feature abstractions, with raw video sometimes omitted or only partially included.

**Figure 6.**
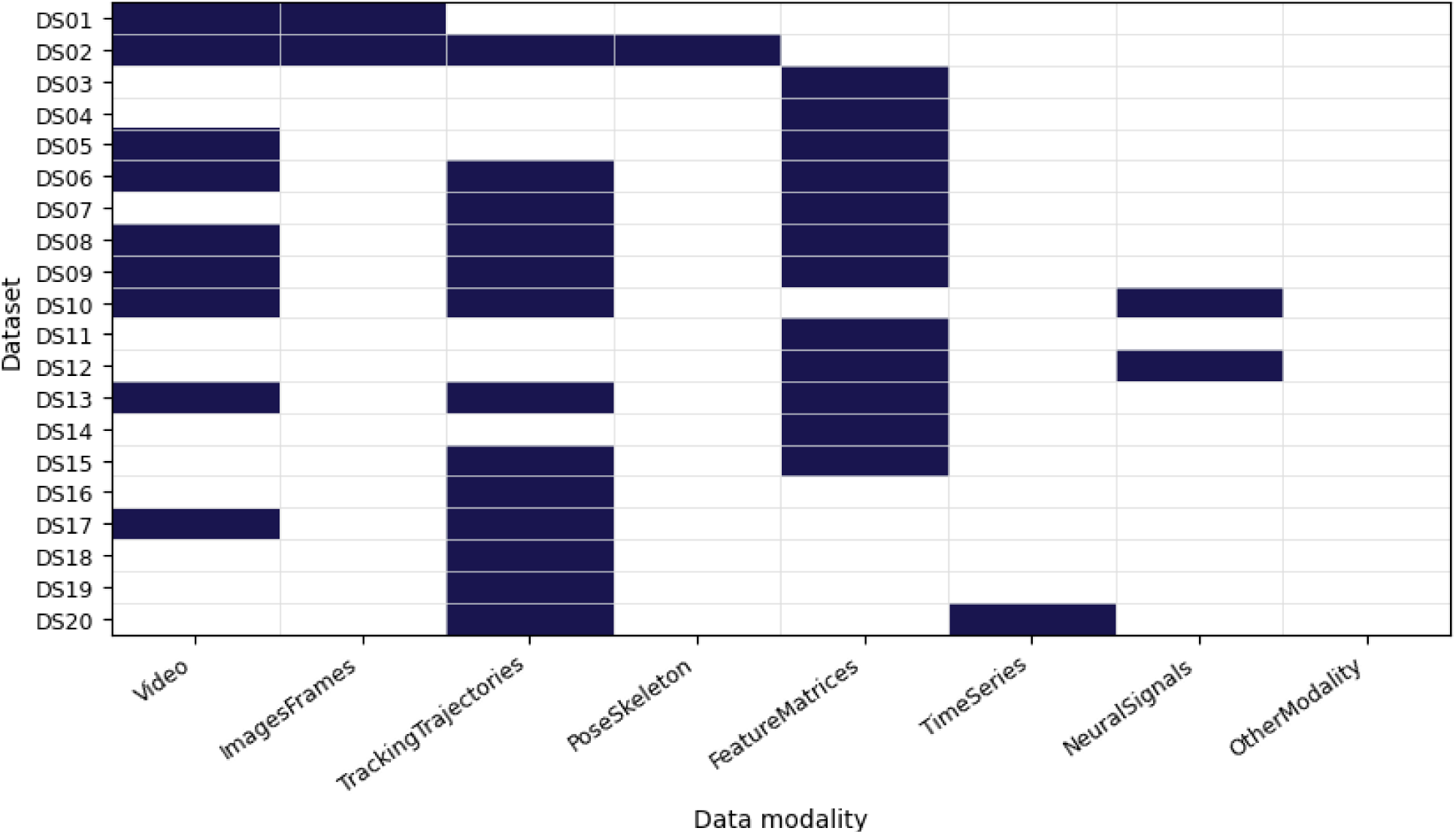
Stage 2 modalities matrix (datasets × modalities).

Open and widely supported formats dominated the corpus (Figure 7). Spreadsheet/tabular files (CSV/TSV/XLSX) appeared in 11 datasets and standard video containers (MP4/AVI/MOV) in 7 datasets. Text-based outputs (TXT/DLM) occurred in 6 datasets, while analysis-ready numeric containers (NPY/PKL/ARFF) appeared in 5 datasets. Legacy or proprietary formats were comparatively uncommon (MAT in 4 datasets; BIN/IPF in a single deposit). Raw–derived separation and provenance ranged from explicit directory-level schemas with clear naming conventions to ad hoc deposits with limited traceability, with the best-performing datasets combining structured directory hierarchies and file naming with explicit links between primary files and derivatives.

**Figure 7.**
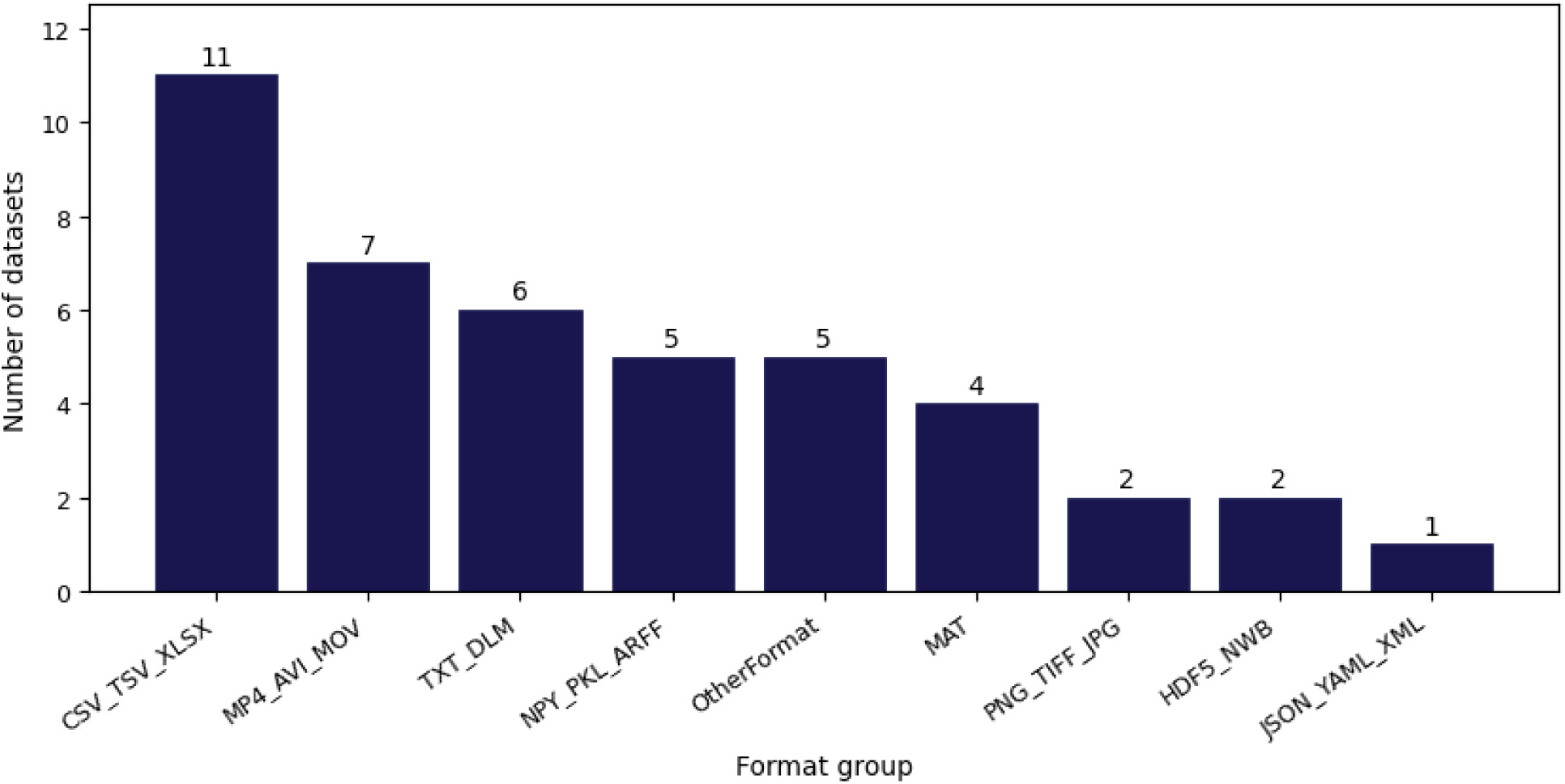
File format distribution across Stage 2 deposits (open vs proprietary/legacy).

##### RQ2 (metadata completeness)

Human-readable documentation was common across deposits, typically including at least a minimal description of organism, experimental task, and acquisition setup. However, machine-readable metadata, structured variable dictionaries, and explicit raw-to-derived provenance were inconsistently provided. Typical gaps included incomplete organism descriptors (e.g. missing strain or developmental stage), narrative-only reporting of experimental conditions, and weak linkage between primary recordings and derived outputs. Only a minority of datasets approached the level of structured metadata seen in curated zebrafish resources.

##### RQ3 (dissemination and reuse)

Deposits were hosted across Zenodo, Dryad, Dataverse, Figshare, and project-specific platforms. All included datasets provided stable landing pages with DOIs or other persistent identifiers (20/20), supporting long-term Findability. Licensing clarity was high overall (19/20 with explicit open licenses); one deposit lacked a dataset-level license statement and was therefore only partially reusable. These dissemination profiles indicate that persistent identifiers and open licensing are now common practice for technically mature invertebrate datasets, even though metadata and standards adoption lag behind.

##### RQ4 (FAIR indicators)

Findability and Accessibility were uniformly satisfied (100% of datasets), reflecting stable PIDs and open repository landing pages (Figure 8). Interoperability was weaker (70%), primarily due to sparse structured metadata, limited standard vocabularies, and minimal use of self-describing formats. Reusability was high (97.5%), driven by near-universal open licensing; the single non-licensed dataset was only partially reusable. Overall, the Stage 2 corpus demonstrates that FAIR-aligned dissemination is achievable in invertebrate behavioural research, but that Interoperability remains the principal bottleneck—especially for AI benchmarking and cross-dataset synthesis.

**Figure 8.**
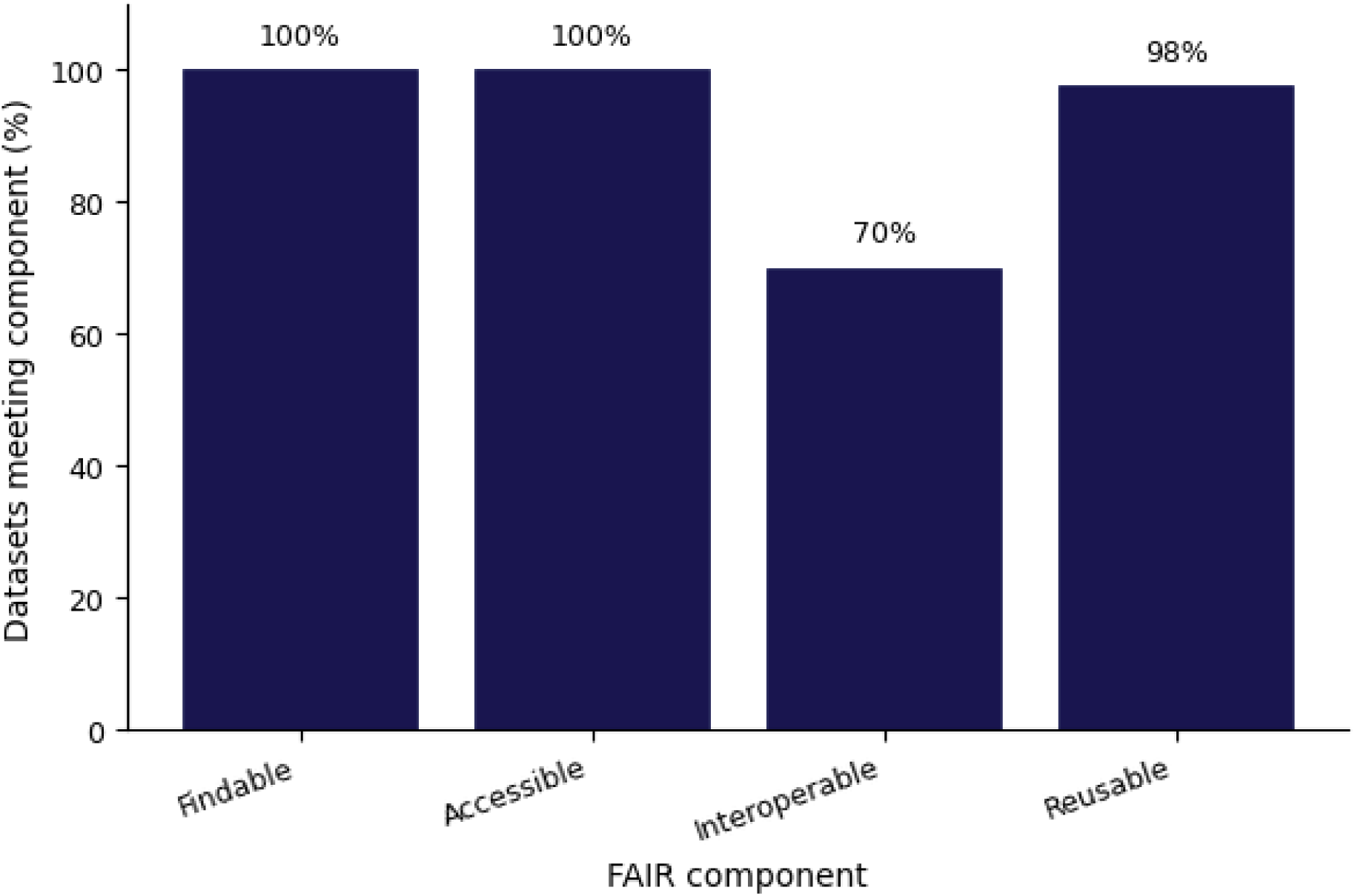
Summary of FAIR indicators across Stage 2 datasets (F/A/I/R).

##### RQ5 (standards/ontologies)

Formal standards and ontology stacks were rare outside curated zebrafish-linked resources. Datasets scoring highest on interoperability consistently separated raw and derived layers with clear traceability, provided machine-readable codebooks, and relied on open, self-describing containers. Most invertebrate deposits remained project-specific in their labelling and did not expose explicit ontology mappings, underscoring both the feasibility and the current scarcity of harmonised behavioural data standards for these models.

#### Technical score distributions and inter-rater agreement

Across dimensions, scores clustered around a median of 2.0 (Figure 9), indicating that most datasets were reusable after moderate preprocessing. Technical quality showed the highest mean score, while usability, annotation richness, and AI-readiness were slightly lower but similar. This pattern suggests that file-level integrity and basic organisation are often stronger than metadata depth or explicit AI-oriented labelling. In other words, many datasets are technically sound but not yet optimised for direct, large-scale reuse in machine-learning pipelines.

**Figure 9.**
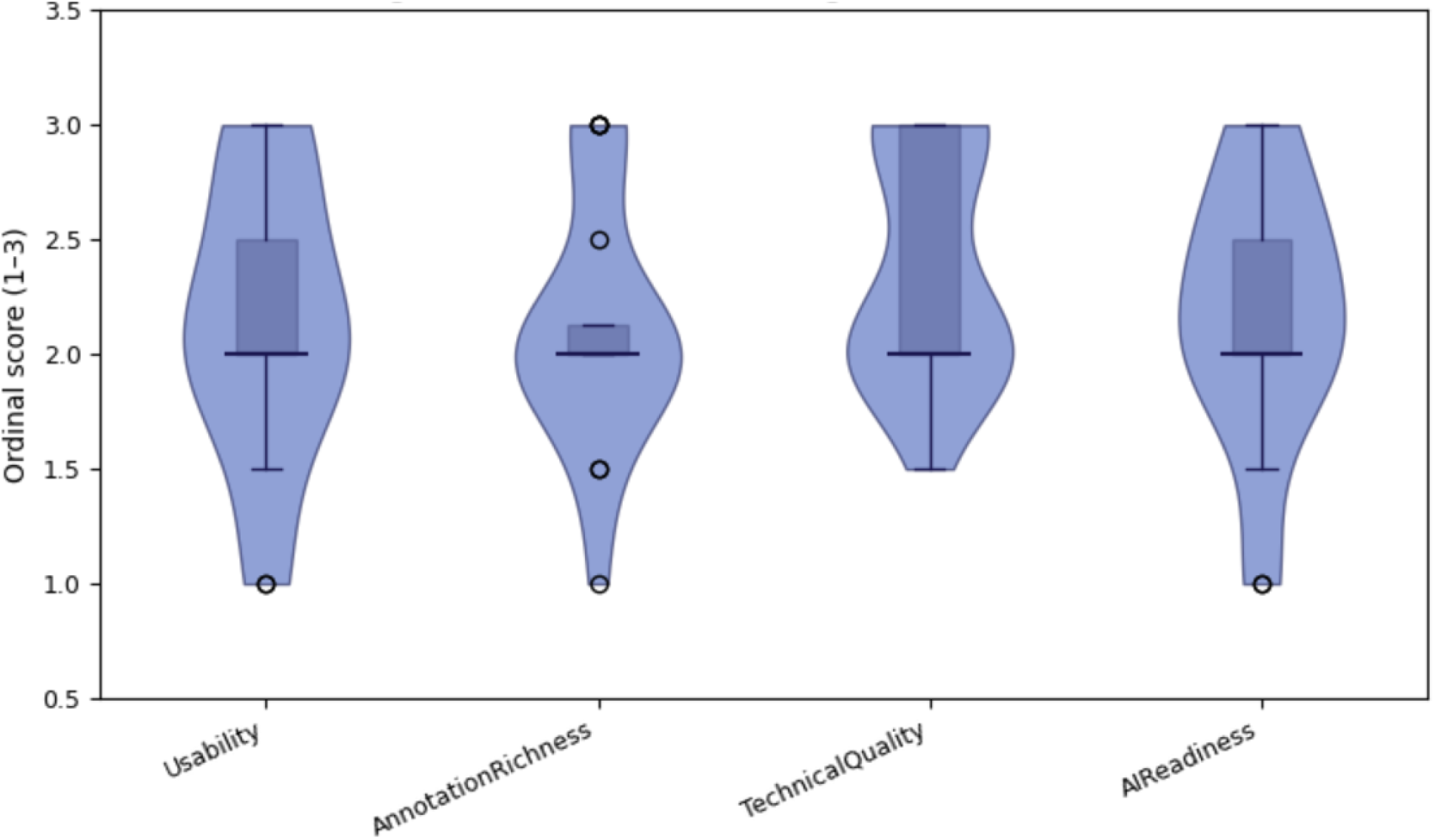
Distributions of Stage 2 technical scores (Usability, Annotation richness, Technical quality, AI-readiness).

Inter-rater agreement (Cohen’s κ) was interpretable for a 1–3 ordinal scale; per-deposit κ values and rater-specific scores are provided in Supplementary Table S9. Lower κ values tended to occur for borderline mixed profiles and reflect the conservative behaviour of κ on three-level ordinal scales. Despite this, most datasets showed at least slight-to-moderate agreement, supporting the robustness of the four-dimension rubric for capturing reuse-relevant differences between deposits. To our knowledge, this is the first systematic, dataset-centred quantification of FAIR indicators and AI-readiness across invertebrate behavioural resources.

### 4. Technical Validation

Technical validation evaluates the reliability and reproducibility of the Stage 2 screening and scoring pipeline rather than experimental findings of the primary datasets. The four-dimension framework—usability, annotation richness, technical quality, and AI-readiness—was designed to reflect core FAIR-oriented reuse enablers as outlined in the FAIR principles[8] and to highlight machine-actionable features relevant for computational and AI workflows.

#### Rationale for the four-dimension scoring system

The rubric captures complementary dimensions relevant to RQ1–RQ4. Ordinal 1–3 thresholds were selected to foreground practical reuse criteria: dataset interpretability, availability of training-ready labels, technical integrity across file layers, and the presence of structured metadata enabling machine actionability. These dimensions extend conventional FAIR alignment by explicitly operationalising the requirements for supervised and unsupervised AI applications.

#### Inter-rater agreement

Two reviewers independently scored all datasets using masked repository identifiers. Inter-rater agreement was quantified using Cohen’s κ[13], a widely used metric for categorical concordance. κ values ranged from −0.33 to 1.00, with most datasets showing slight-to-moderate agreement. Interpretation followed the Landis–Koch thresholds[14], according to which κ < 0.00 indicates poor agreement, 0.00–0.20 slight, 0.21–0.40 fair, 0.41–0.60 moderate, and ≥0.61 substantial to almost perfect agreement. Lower κ values typically reflected borderline mixed profiles and the conservative behaviour of κ on three-level ordinal scales. Rater-level scoring sheets and κ values are released in Supplementary Table S9 and Supplementary Data File S1, allowing independent recalculation or alternative aggregation schemes.

#### Reproducibility checks

During screening, all datasets underwent systematic verification of raw–derived linkage, internal integrity, and basic executability of included scripts or workflows (where applicable). All judgments were recorded in standardised extraction templates and reconciled after re-inspection. This allowed identification of inconsistencies in file structures, undocumented preprocessing steps, outdated software dependencies, and potential reusability barriers.

#### Robustness

Sensitivity analyses examined whether excluding the most problematic datasets—those with irrecoverable broken links, non-functional code, or severely incomplete metadata—altered downstream conclusions. Exclusion did not meaningfully change the observed patterns in modality prevalence, dominance of open formats, FAIR-alignment deficits, or AI-readiness profiles. Thus, the qualitative conclusions reflected in Figures 6–9 were robust to dataset-level variability.

### 5. Usage Notes

The Stage 1 and Stage 2 catalogue is intended to support reuse of invertebrate behavioural datasets in three main contexts: secondary biological analyses, FAIR-oriented auditing, and benchmarking of AI methods. Users are encouraged to begin by defining a concrete reuse objective and then narrowing candidate datasets using the Stage 2 descriptors (organism, modality, file formats, provenance, licensing, and technical score profiles). In practice, these descriptors allow rapid triage of deposits that are likely to be immediately actionable versus those requiring extensive preprocessing or metadata reconstruction. Interpretation guidelines for the four technical dimensions by typical use case are summarised in Table 3.

**Table 3.** Interpreting Stage 2 technical scores by use case.

| Use case | Recommended score profile | Minimal acceptable thresholds | Typical limiting factors if below threshold |
| --- | --- | --- | --- |
| Biological reanalysis (hypothesis testing, replication, meta-analysis) | Balanced profile, strongest weight on Usability + Technical quality; Annotation richness optional unless behaviours are categorical | Usability $\geq 2$, Technical quality $\geq 2$ | unclear organisation, missing provenance, narrative-only metadata |
| Supervised AI (classification, detection, pose-to-behaviour models) | High Annotation richness + AI-readiness, supported by strong Technical quality | Annotation richness $\geq 2$, AI-readiness $\geq 2$, Technical quality $\geq 2$ | absent/weak labels, incomplete codebooks, no raw-derived link |
| Unsupervised AI (clustering, representation learning, anomaly detection) | High Technical quality + Usability; labels not required if derived trajectories/features are consistent | Technical quality $\geq 2$, Usability $\geq 2$ | format heterogeneity, missing variable definitions, unstable access |

A recurring decision in reuse workflows concerns whether to operate on raw recordings or derived layers. Many deposits provide both primary video/imaging and processed tracking or feature outputs, but their traceability is not always explicit. For biological reanalysis, derived layers may be sufficient when behavioural variables are clearly defined and adequately documented. For AI applications, the choice depends on task type: supervised modelling requires raw data paired with training-ready labels and explicit raw–derived linkage, whereas unsupervised approaches can often rely on high-quality trajectories or feature matrices alone. The four Stage 2 technical scores should therefore be interpreted as a joint profile rather than in isolation. For example, a dataset can be technically robust and usable, yet still unsuitable for supervised learning if behavioural labels or codebooks are absent; conversely, rich annotation without clear organisation may still be valuable but implies a higher reuse cost.

Across Stage 2 deposits, common reuse caveats include missing persistent identifiers in some GitHub-only resources, inactive third-party links, occasional proprietary or legacy formats, and weak provenance between primary recordings and derived outputs. Another practical limitation is that some datasets depend on older software environments, which may require containerisation or manual updating for reproducible execution. These issues are documented per dataset in Supplementary Table S9 and should be considered when estimating feasibility and time required for reuse.

Despite these limitations, many deposits can be substantially improved through post hoc harmonisation. Because custom folder structures and narrative-only metadata are frequent in invertebrate behavioural resources, downstream users can often increase interoperability by refactoring documentation into machine-readable codebooks, standardising variable names and units, mapping behavioural labels to community ontologies (e.g., NBO or ZECO/OBI), and packaging deposits into self-describing hierarchies such as NWB/HDF5-like containers or RO-Crate. Such steps typically strengthen Interoperability and AI-readiness without modifying the primary data, enabling more robust cross-dataset synthesis, transfer learning, and computational benchmarking.

Looking ahead, the Stage 1 and Stage 2 catalogue also exposes a broader gap in the current landscape of behavioural data sharing: technically mature, openly documented datasets are heavily concentrated in a small group of canonical model organisms. Extending similar FAIR- and AI-oriented practices to a wider range of invertebrate and non-model taxa will be essential for capturing the full diversity of animal behaviour, enabling more generalisable AI benchmarks, and avoiding model-centric biases in downstream biological inference.

### 6. Limitations

Despite broad PRISMA-guided searching and an independent multi-repository screening workflow, some relevant datasets may have been missed. This risk is highest for deposits lacking descriptor publications, using sparse or inconsistent metadata, or being weakly indexed within repository search engines. Discovery sensitivity differed across platforms because several specialised archives did not provide fully functional dataset-level search interfaces during the screening window, limiting systematic querying.

Although the Stage 2 ordinal rubric was pre-thresholded, piloted, and independently applied by two reviewers, it necessarily involves qualitative judgement. Inter-rater agreement should therefore be interpreted conservatively, especially given the tendency of Cohen’s κ to underestimate agreement on three-level ordinal scales and in borderline mixed-quality profiles. Users combining or re-weighting the four technical dimensions for specific applications should consider re-deriving summary scores from the released rater-level sheets.

Finally, the survey focused on major invertebrate models with larval zebrafish as a structured comparator. The observed dissemination, FAIR-alignment, and AI-readiness patterns may not fully generalise to less-established taxa or behavioural domains not represented in the included corpus. At the same time, the concentration of technically well-described, openly shared behavioural datasets in a small subset of model organisms is itself an important finding, highlighting a structural gap in data availability for many invertebrate and non-model taxa.

## Supporting information

Supplementary Table S1. Full database-specific search queries, filters, run dates, and retrieved counts. (2)

Supplementary Table S4. Standards and ontologies adoption (RQ5). (1)

Supplementary Table S5b. Decision thresholds for overall record quality _ risk of reporting-accessibility bias (Stage 1). (1)

Supplementary Table S6. Stage 2 repository search strategy_ keyword families and thematic query blocks. (1)

Supplementary Table S7. Stage 2 repository search log (1)

Supplementary Table S9. Stage 2 ordinal scores and inter-rater agreement per dataset. (1)

Supplementary Table S11. Source-resolved PRISMA 2020 flow summary for Stage 1.

## Data Availability

All data generated for this survey—including Stage 1 PRISMA extraction tables, Stage 2 repository search logs, the Stage 2 master extraction sheet, rater-level scoring and κ estimates, and derived summary statistics—are available as Supplementary Information accompanying this Data Descriptor (Supplementary Tables S1–S10 and Supplementary Data Files S1 and S2). No additional datasets were deposited in an external public repository; all figures and summary results can be regenerated directly from these Supplementary Tables and Data Files.

## Code Disclosure

Custom code is not publicly released because it consisted solely of simple, dataset-specific helper scripts used to download, open, and run individual datasets (basic file loading, format conversion, execution checks) without implementing novel analytical methods. All intermediate and final products required to reproduce the analyses and figures are contained in the released Supplementary Tables and Data Files; thus, all figures can be regenerated directly from the publicly available extraction sheets without access to custom scripts. Where relevant, repository-linked code notebooks or pipelines provided by original dataset authors are documented in the Stage 2 descriptors to support further reuse.

## 7. Supplementary Materials

1. Supplementary Table S1. Full database-specific search queries, filters, run dates, and retrieved counts.
2. Supplementary Table S2. Dataset technical characteristics (RQ1–RQ3).
3. Supplementary Table S3. FAIR dissemination and compliance (RQ3–RQ4).
4. Supplementary Table S4. Standards and ontologies adoption (RQ5).
5. Supplementary Table S5. Reporting/accessibility bias(Stage 1).
6. Supplementary Table S5a. Record-level scoring rubric for reporting/accessibility bias (Stage 1).
7. Supplementary Table S5b. Decision thresholds for overall record quality / risk of reporting-accessibility bias (Stage 1).
8. Supplementary Table S6. Repository search strategy: keyword families and thematic query blocks (Stage 2).
9. Supplementary Table S7. Repository search log (Stage 2).
10. Supplementary Table S8. Definitions and ordinal scoring thresholds (1–3) for Stage 2 technical quality dimensions.
11. Supplementary Table S9. Stage 2 ordinal scores and inter-rater agreement per dataset.
12. Supplementary Table S10. Data extraction template and coding scheme for dataset evaluation (RQ1–RQ4).
13. Supplementary Table S11. Source-resolved PRISMA 2020 flow summary for Stage 1.

## Supplementary Data Files

Supplementary Data File S1. Stage 2 master extraction sheet (CSV/XLSX). Supplementary Data File S2. Full repository search audit trail for Stage 2 (CSV).

## Notes

### Competing Interest Statement

The authors have declared no competing interest.

