## Supplementary Table S1. Full database-specific search queries, filters, run dates, and retrieved counts. (2) for "Systematic Survey of Public Datasets for Behavioral Research in Invertebrate Models: Toward FAIR and Standardized Data Sharing"

| Database / Platform | Interface / Provider | Search date | Time window | Full search string (as executed) | Search fields | Filters / Limits applied | Records retrieved (n) | Notes |
|---|---|---|---|---|---|---|---|---|
| PubMed/ MEDLINE | PubMed Advanced Search | 2025-06-30 | 2015-01-01–2025-06-30 | (("behavior*" OR "behaviour*" OR locomotion OR "motion tracking" OR ethogram OR "pose estimation") AND (dataset* OR "open data" OR repository OR "data descriptor") AND ("Drosophila melanogaster" OR Drosophila OR "Caenorhabditis elegans" OR C. elegans OR "Galleria mellonella" OR planarian* OR "Schmidtea mediterranea" OR "Danio rerio" OR zebrafish)) | Title/Abstract | English; Journal Article | **n = 283** | Manual exclusion during screening: review papers removed only if they did not provide dataset links (criterion applied post-search, not a platform filter). Exported as .nbib. |
| Scopus | Elsevier Scopus | 2025-06-28 | 2015-01-01–2025-06-30 | TITLE-ABS-KEY ( (behavior* OR behaviour* OR locomotion OR "motion tracking" OR ethogram OR "pose estimation") AND (dataset* OR "open data" OR repository OR "data descriptor") AND ("Drosophila melanogaster" OR drosophila OR "Caenorhabditis elegans" OR "Galleria mellonella" OR "Schmidtea mediterranea" OR planarian* OR "Danio rerio" OR zebrafish) ) | Title/Abstract/ Keywords | English; Document type: Article, Conference Paper; Subject areas: Neuroscience, Bioinformatics, CS | **n = 254** | Subject-area filter used to reduce off-topic retrieval while maintaining sensitivity, based on pilot runs; complemented by broader non-indexed screening. Exported RIS. |
| Web of Science Core Collection | Clarivate WoS | 2025-06-27 | 2015-01-01–2025-06-30 | TS = ((behavior* OR behaviour* OR locomotion OR "motion tracking" OR ethogram OR "pose estimation") AND (dataset* OR "open data" OR "data repository" OR "data sharing" OR metadata OR "data descriptor" OR "data paper") AND ("Drosophila melanogaster" OR Drosophila OR "Caenorhabditis elegans" OR "C. elegans" OR "Galleria mellonella" OR planarian* OR "Schmidtea mediterranea" OR "Danio rerio" OR zebrafish OR invertebrate*) AND (FAIR OR ontology OR "data standard*" OR "data descriptor" OR "data paper")) | Topic | English; Article/Proceedings; Timespan limited to window | **n = 805** | Corrected for reproducibility: added parentheses to preserve Boolean logic and removed duplicated dataset block present in draft; conceptual structure aligned with other databases. |
| IEEE Xplore | IEEE Xplore Advanced | 2025-06-25 | 2015-01-01–2025-06-30 | ("behavior" OR "behaviour" OR locomotion OR "pose estimation") AND (dataset OR "open data") AND (Drosophila OR "C. elegans" OR planarian OR zebrafish) | Metadata only | English; Content type: Journals + Conferences | **n = 19** | Query shortened due to 15-term platform cap; highest-recall behavior/data/organism terms retained (e.g., removed lower-yield synonyms such as ethogram/motion tracking). |
| ACM Digital Library | ACM DL Advanced | 2025-06-25 | 2015-01-01–2025-06-30 | ([[All: behavior*] OR [All: behaviour*] OR [All: locomotion] OR [All: ethogram] OR [All: "motion tracking"] OR [All: tracking] OR [All: "pose estimation"] OR [All: "behavioral phenotyping"] OR [All: "behavioural phenotyping"] OR [All: "trajectory tracking"] OR [All: "animal tracking"]]) AND ([[All: dataset*] OR [All: "data set"] OR [All: "open dataset"] OR [All: "public dataset"] OR [All: "benchmark dataset"] OR [All: repository] OR [All: "data descriptor"] OR [All: "data release"] OR [All: "open data"]]) AND ([[All: Drosophila] OR [All: "Drosophila melanogaster"] OR [All: "Caenorhabditis elegans"] OR [All: "C. elegans"] OR [All: planarian] OR [All: planarians] OR [All: "Schmidtea mediterranea"] OR [All: "Galleria mellonella"] OR [All: zebrafish] OR [All: "Danio rerio"]]) AND ([[All: video] OR [All: imaging] OR [All: track*] OR [All: "pose"] OR [All: "behavior recognition"]]) | All fields | English; full-text where available | **n = 159** | Syntax adjusted to ACM field-tag format. |
| Google Scholar | Google Scholar | 2025-06-20 | 2015-01-01–2025-06-30 | "behavior dataset" (Drosophila OR "C. elegans" OR planarian OR zebrafish) tracking | All fields | Year-range limited to window; sorted by relevance; first 200 results screened | **n = 200** | Non-indexed source screened per PRISMA-S practice to reduce indexing/publication bias. |
| bioRxiv (preprints) | bioRxiv search + manual screen | 2025-06-20 to 2025-06-30 | 2015-01-01–2025-06-30 | Manual search using core concepts from indexed strings (behavior/locomotion/pose estimation + dataset/open data + model organisms) | All fields | English; relevance screening of returned preprints | **n = 57** | Added to capture recent datasets/method papers not yet indexed. |
| arXiv (preprints) | arXiv search + manual screen | 2025-06-20 to 2025-06-30 | 2015-01-01–2025-06-30 | Manual search using core concepts from indexed strings (behavior tracking/pose estimation + dataset/open data + model organisms) | All fields | English; relevance screening of returned preprints | **n = 214** | Added to reduce time-lag bias for CS/AI dataset releases. |
| Reference list screening / citation chasing | Backward + forward screening | 2025-06-20 to 2025-06-30 | No restriction beyond main window | Reference lists of all included studies screened; forward citations checked in WoS/GS | N/A | Manual snowballing | **n = 67** | Complements database searches; recommended by PRISMA-S for completeness. |
