## Supplemental Data 1 for "Systematic Survey of Public Datasets for Behavioral Research in Invertebrate Models: Toward FAIR and Standardized Data Sharing"

**Supplementary Table S2**. Dataset technical characteristics (RQ1–RQ3).

| No. | Source (author, year, DOI) | Dataset | Organism | Data type | Data format | Metadata format | Folder structure / file organization | Metadata description | Time stamps (sampling / rate) | Multimedia format | Compliance notes | Standardization notes | Search source |
|---|---|---|---|---|---|---|---|---|---|---|---|---|---|
| 1 | Castro et al., 2023, Heliyon. “Automatic segmentation of Caenorhabditis elegans skeletons…” | Dataset of real and synthetic low-resolution worm aggregations | *C. elegans* | Low-resolution grayscale images; skeleton tracks | PNG (images), PTS (text), XML | XML (skeleton points), PTS | Images with paired .XML/.PTS files organized as time sequences | Skeleton points (X,Y), color, width | Yes; 30 fps, 1 Hz summary | PNG (1944×1944), PTS, XML | Detailed experimental/breeding and imaging description | Well documented, but no reference to external metadata standards | Web of Science |
| 2 | García-Garví et al., 2023. “Automation of C. elegans lifespan assay…” | Synthetic + real dataset for lifespan assay automation | *C. elegans* | Synthetic images, experimental videos, segmentation masks, age/motion classes | PNG, AVI, XML | XML masks; no formal metadata schema | Separate folders for synthetic vs. real; masks separated from originals | Mask positions, age classes, alive/dead states encoded in XML + repository text | Yes; frame every 5 min over ~14 days | PNG (binary/RGB), AVI | Detailed model/training+imaging conditions | Good file structure, but no formal ontology/metadata standard | Web of Science |
| 3 | Layana-Castro et al., 2023, Int. J. Computer Vision. “Skeletonizing C. elegans…” | Synthetic + real dataset of worm aggregations with skeleton labels | *C. elegans* | Images + skeleton segmentation | PNG, XML, JSON | XML/JSON (points, masks, classes) | Grouped by experiments; synthetic/real separated; masks separate | Skeleton points (x,y), IDs, widths, binary masks | Yes; 30 fps (real sequences) | PNG, JSON, XML | Thorough synthetic-data pipeline description | No external FAIR/ontology alignment reported | Web of Science |
| 4 | Bradford et al., 2017. “Zebrafish Models of Human Disease at ZFIN” | ZFIN (Zebrafish Information Network) | *Danio rerio* | Genotypes, phenotypes, behavioral annotations, gene–disease links | Online relational DB; ZDB IDs | ZFA, PATO, ZECO, HPO, MP (ontologies) | Relational structure; entity links via ZDB identifiers | Standardized ontology-based annotations | NA (not time-sampled) | Primarily symbolic/textual; some phenotypic media | Curated by ZFIN with explicit provenance links | High compliance; ontology-driven interoperability | Scopus |
| 5 | Ruzicka et al., 2015, DOI:10.1002/dvg.22868 | ZFIN database update | *Danio rerio* | Genotypes, phenotypes, expression, disease models | Relational DB; TSV/TXT, JSON, RDF exports | ZFA, ZECO, GO, PATO, HPO, MP, mCODE | Web interface + downloadable files + REST | Ontology annotations linked to experiments/publication s | NA (descriptive) | No native video datasets | Extensive provenance and stable access | Model FAIR biological DB; strong ontology support | Google Scholar |
| 6 | Pham et al., 2023. “Eigenfeature-Enhanced LSTM…” | EigenWorm posture dataset (Stephens et al., MBL) | *C. elegans* | Postural trajectories; PCA components; behavior labels | CSV, JSON | No formal schema; narrative only | Processed files only; no official directory in paper | PCA (6 eigenworms), behavior labels, durations | Yes; 2-s samples | None (vector data only) | Data provenance described in paper | No FAIR metadata/ontology alignment | Google Scholar |
| 7 | Liu et al., 2015, DOI:10.1371/journal.pone.0139521 | Zebrafish VMR dataset (Harvard Dataverse) | *Danio rerio larvae (5 dpf)* | Locomotor activity time series under light stimulation | CSV, TXT | Repository narrative; no schema | Experimental folders by condition/plate | Movement parameters, IDs, cycles | Yes; 1 Hz for 1800 s | None | Detailed experiment protocol | No standards/ontologies reported | Google Scholar |
| 8 | Štih et al., 2019. “Stytra…” | Stytra example datasets (GitHub/Zenodo) | *Danio rerio larvae (5–7 dpf)* | Trajectories, stimulation logs, closed-loop data | CSV, HDF5, MP4 | YAML + JSON protocol description | Structured folders by experiment/session/fish | YAML docs + stimulus/behavior logs | Yes; real-time stamps + sync | MP4 + coordinate CSV | Complete open experimental system | Very high reuse readiness; FAIR-friendly formats | Google Scholar |
| 9 | Hsieh et al., 2023, DOI:10.3390/toxics11050407 | SEAZIT interlaboratory toxicity dataset | *Danio rerio embryos/larvae* | Phenotypic observations; binary + continuous outcomes | CSV, XLSX | Not reported | Tabular files grouped by variables/outcomes | Phenotype variable definitions in text | Yes; selected exposure time points | None | Detailed protocol and inter-lab standardization | No semantic/FAIR schemas reported | Scopus |
| 10 | Johnson et al., 2020, DOI:10.1016/j.cub.2019.11.026 | Tail kinematics + behavioral states (Dryad) | *Danio rerio larvae (5–7 dpf)* | Tail kinematics, latent state models, sequences | HDF5, CSV | README + notebooks | Structured folders by session/larva/model | Variables + protocols in README/notebooks | Yes; 700 Hz sampling | HDF5 time-series; some media | Very detailed protocol + code | No formal ontologies (e.g., ZECO/NBO) | Google Scholar |
| 11 | Gendelev et al., 2024. “Deep phenotypic profiling…” | DeepFish behavioral dataset | *Danio rerio larvae (6–7 dpf)* | Motion index time series from 3 screens | .npy, CSV | README + metadata CSV | Grouped by screen/plate; ID mapping provided | Chemical name, MI, plate info, labels | Yes; MI sampled every 1 s–1 min | None (metric only) | Highly repeatable screen with QC | No behavioral ontologies reported | Web of Science |
| 12 | Van Slyke, 2018, DOI:10.1007/978-1-4939-7737-6_11 | ZFIN usage chapter | *Danio rerio* | Broad ZFIN data types incl. limited behavior | HTML interface + TSV/XML/OBO/RDF exports | ZECO, PATO, ZFA, ZFS, GO, NBO, OBI etc. | Standard ZFIN object model | Ontology-based metadata; stable IDs | Yes; developmental stage/time refs | Links to images/videos embedded | Comprehensive FAIR model DB | Fully ontology-aligned | Google Scholar |
