## Supplemental Data 2 for "Systematic Survey of Public Datasets for Behavioral Research in Invertebrate Models: Toward FAIR and Standardized Data Sharing"

**Supplementary Table S3**. FAIR dissemination and compliance (RQ3–RQ4).

| No. | Dataset name | Organism | Sharing platform | License | Data format | Access method (link/API) | FAIR assessment (F/A/I/R) | FAIR-related notes |
|---|---|---|---|---|---|---|---|---|
| 1 | Synthetic + real C. elegans skeleton dataset | *C. elegans* | GitHub + ZIP | CC BY-NC-ND stated in paper; no dataset-level license | PNG, PTS, XML | Direct download; no API | Partially; I limited | No persistent identifier; no interoperable metadata (no RO-Crate/JSON-LD; no ontologies). |
| 2 | C. elegans lifespan assay automation dataset | *C. elegans* | GitHub + ZIP (Active Vision) | Not reported (open repository only) | PNG, AVI, XML | Direct download | Partially; I limited | No DOI; license unclear; no machine-readable metadata/ontologies. |
| 3 | C. elegans skeletonization dataset | *C. elegans* | GitHub + download link | Not reported (default GitHub) | PNG, XML, JSON | Direct download | Partially; I limited | No DOI/DataCite record; no JSON-LD/RDF/RO-Crate. |
| 4 | ZFIN | *Danio rerio* | ZFIN.org | CC BY 4.0 | Relational DB; RDF/JSON/TSV exports | Web + REST API | Comprehensively F/A/I/R | Model FAIR organism DB with maintained versions and rich ontology coverage. |
| 5 | ZFIN (updates & directions) | *Danio rerio* | ZFIN.org | CC BY 4.0 | JSON, RDF, TSV | Web + REST API | Comprehensively F/A/I/R | Strong PID, licensing, API, and ontology interoperability. |
| 6 | EigenWorm dataset | *C. elegans* | wormbehavior.mbl.edu | Not reported | CSV, JSON | Direct download | Partially; I limited | No DOI; no formal schema; no ontologies. |
| 7 | Zebrafish VMR dataset | *Danio rerio* | Harvard Dataverse | CC0/CC-BY default in Dataverse | CSV, TXT | DOI link; no API | Partially; I limited | Good Findable/Accessible; limited interoperability (no ontologies/JSON-LD). |
| 8 | Stytra example datasets | *Danio rerio* | GitHub + Zenodo | GPLv3 (software), CC-BY (data) | CSV, HDF5, MP4, JSON | DOI + repo; export tools | Comprehensively F/A/I/R | Open formats, rich metadata, stable versions, strong reuse support. |
| 9 | SEAZIT toxicity dataset | *Danio rerio* | US EPA HERO / on request | Not reported | CSV, XLSX | No DOI; no API | Partially; I limited | Open in practice but no persistent ID, license statement, or interoperable metadata. |
| 10 | Zebrafish tail kinematics dataset | *Danio rerio* | Dryad + GitHub | CC0 | HDF5, CSV | DOI + notebooks | Partially; I limited | Strong F/A; interoperability limited by lack of formal ontologies. |
| 11 | DeepFish behavioral dataset | *Danio rerio* | Zenodo + GitHub | CC-BY 4.0 | .npy, CSV, JSON | DOI + repository | Partially; I limited | Well documented; lacks RO-Crate/JSON-LD and ontology mapping. |
| 12 | ZFIN (usage chapter) | *Danio rerio* | ZFIN.org | Custom open reuse w/ citation | TSV/XML/OBO/RDF exports | Web + REST API | Comprehensively F/A/I/R | Prime FAIR/ontology-aligned reference repository. |
