## Supplementary Table S4. Standards and ontologies adoption (RQ5). (1) for "Systematic Survey of Public Datasets for Behavioral Research in Invertebrate Models: Toward FAIR and Standardized Data Sharing"

| No. | Use case (citation / DOI) | Standard / ontology name | Type | Source / initiative | Application area | FAIR relevance | Evidence of use in dataset | Adoption level | Notes |
|---|---|---|---|---|---|---|---|---|---|
| 1 | Castro et al., 2023 (Heliyon) | Not reported | NA | NA | NA | Partially (I limited) | No | NA | No reference to NBO/OBI/BCO or metadata schemas. |
| 2 | García-Garví et al., 2023 | Not reported | NA | NA | NA | Partially (I limited) | No | NA | Behavior classes not mapped to formal ontologies. |
| 3 | Layana-Castro et al., 2023 (IJCV) | Not reported | NA | NA | NA | Partially (I limited) | No | NA | Dataset uses custom labels only. |
| 4 | Bradford et al., 2017 (ZFIN) | ZFA; ZECO; HPO; MP; PATO | Ontologies | OBO Foundry / Monarch / MGI / ZFIN | Anatomy, experimental conditions, phenotypes, cross-species disease/behavior links | Comprehensive | Yes | Widespread / growing | Core ontology stack for zebrafish DB interoperability. |
| 5 | Ruzicka et al., 2015 (ZFIN) | ZFA; ZECO; GO; HPO; MP; PATO; mCODE | Ontologies | OBO / GO Consortium / Monarch / MGI / ZFIN | Anatomy, conditions, gene function, phenotypes, disease links | Comprehensive | Yes | Widespread | Stable standards-based architecture. |
| 6 | Pham et al., 2023 (EigenWorm) | Not reported | NA | NA | Behavior labels (forward/reversal/pause) | Partially (I limited) | No | NA | Labels are informal; no ontology cross-walk. |
| 7 | Liu et al., 2015 (VMR) | Not reported | NA | NA | Locomotion metrics | Partially (I limited) | No | NA | No ZECO/NBO/OBI/UO or schema.org. |
| 8 | Štih et al., 2019 (Stytra) | Custom YAML/JSON schema (no formal ontology) | Data schema | Stytra / Portugues Lab | Stimulation, tracking, closed-loop behavior | Comprehensive | Yes | Growing | Highly structured; ontology mapping feasible but not implemented. |
| 9 | Hsieh et al., 2023 (SEAZIT) | Not reported | NA | NA | Toxicology phenotypes | Partially (I limited) | No | NA | Custom textual phenotype nomenclature only. |
| 10 | Johnson et al., 2020 (tail kinematics) | Not reported | NA | NA | Kinematics + latent states | Partially (I limited) | No | NA | No formal behavioral ontology use despite strong structure. |
| 11 | Gendelev et al., 2024 (DeepFish) | Custom schema (no formal ontology) | Data schema | Berger Lab / DeepFish pipeline | High-throughput behavioral phenotyping | Partially (I limited) | No | Growing | Structured labels; no ZECO/NBO/OBI mapping. |
| 12 | Van Slyke, 2018 (ZFIN) | ZFA; ZECO; GO; HPO; MP; PATO; NBO; OBI etc. | Ontologies | OBO Foundry / Monarch / MGI / ZFIN | Anatomy, conditions, phenotypes, behavior links | Comprehensive | Yes | Widespread | Reference standard stack for model-organism data. |
