## Supplemental Data 3 for "Systematic Survey of Public Datasets for Behavioral Research in Invertebrate Models: Toward FAIR and Standardized Data Sharing"

**Supplementary Table S5a.** Record-level scoring rubric for reporting/accessibility bias (Stage 1). Points rubric (0–2) across three dimensions.

| Dimension | Score 2 (High) | Score 1 (Moderate) | Score 0 (Low) |
|---|---|---|---|
| **Documentation completeness** | Full technical description of the dataset, including modalities, file formats, internal structure, raw↔derived linkage, and a machine-readable README/variable dictionary (or equivalent). | Partial description: modalities + repository/persistent link reported, and at least one additional technical element (e.g., file format or variable/field overview), but missing full structure/metadata detail. | Fragmentary description or no meaningful dataset description. |
| **Metadata / FAIR / licensing transparency** | Persistent identifier (DOI/accession), explicit license, and clearly reported FAIR-relevant elements/metadata fields enabling verification and reuse. | Persistent link/identifier and some metadata reported, but license unclear and/or FAIR elements only vaguely mentioned. | No verifiable persistent link/DOI, no metadata beyond narrative, and no license statement |
| **Standards / ontologies reporting** | Standard(s)/ontology explicitly named and its concrete implementation in the dataset described (e.g., annotation schema, controlled vocabulary fields, or file-format standard). | General mention of standards/ontologies without evidence of implementation in the dataset. | No mention, or mention unrelated to the dataset. |
