## Supplementary Table S5b. Decision thresholds for overall record quality _ risk of reporting-accessibility bias (Stage 1). (1) for "Systematic Survey of Public Datasets for Behavioral Research in Invertebrate Models: Toward FAIR and Standardized Data Sharing"

**Supplementary Table S5b.** Decision thresholds for overall record quality / risk of reporting-accessibility bias (Stage 1). Overall classification based on the summed three-dimension score (0–2 points each; total 0–6).

| Total score (0–6) | Overall record quality / risk |
|---|---|
| 5–6 | High (low risk) |
| 3–4 | Moderate (medium risk) |
| 0–2 | Low (high risk) |
