## Supplementary Table S6. Stage 2 repository search strategy_ keyword families and thematic query blocks. (1) for "Systematic Survey of Public Datasets for Behavioral Research in Invertebrate Models: Toward FAIR and Standardized Data Sharing"

| Query family / theme | Organism terms (examples) | Modality / behavior terms (examples) | FAIR / accessibility / ML terms (examples) | Dataset-type terms (examples) | Example combined query |
|---|---|---|---|---|---|
| Larvae – video – tracking | larvae, “insect larva”, zebrafish larvae | video, tracking, tracked, locomotion, movement, behavio | annotation, machine learning (when relevant) | “video dataset”, “open dataset” | (“larvae” OR “insect larva”) AND (tracking OR annotation) AND video |
| Invertebrate FAIR / ML datasets (broad) | invertebrate(s), model organism(s) | behavior, behavioral classification, tracking, pose / movement | “machine learning”, multimodal, FAIR, “open dataset” | “behavioral dataset”, “video dataset”, “tracking dataset” | invertebrate AND (tracking OR annotation OR “open dataset”) AND (behavior OR video) |
| Model-organism generic datasets | “model organism”, invertebrate, Drosophila, “C. elegans”, “Galleria mellonella”, zebrafish | behavior, movement, trackin | “machine learning”, multimoda | “video dataset”, “tracking dataset”, “annotation dataset” | “model organism” AND (video dataset OR tracking dataset OR annotation dataset) |
| Organism-specific deep dives | Drosophila, “C. elegans”, “Caenorhabditis elegans”, “Galleria mellonella”, zebrafish, “Danio rerio” | behavior, movement, tracking, locomotion, annotation | “open dataset”, “behavior classification”, multimodal | dataset, “video dataset” | (Drosophila OR “C. elegans” OR “Galleria mellonella” OR zebrafish) AND (movement OR tracking OR behavior) |
