## Supplementary Table S7. Stage 2 repository search log (1) for "Systematic Survey of Public Datasets for Behavioral Research in Invertebrate Models: Toward FAIR and Standardized Data Sharing"

| Search round (theme) | Date range (execution) | Repositories searched | Example query family | Total results screened | Total included | Notes |
|---|---|---|---|---|---|---|
| Round 1: larvae/video/tracking | 04–23.07.2025 | Google Dataset Search, Zenodo, Dryad, Harvard Dataverse, Figshare, IEEE DataPort, OSF | (“larvae” OR “insect larva”) AND (tracking OR annotation) AND video | 26 | 7 | NWB-aligned platforms, Behavioral Data Commons, and DANDI had no functional dataset search engines at search time. |
| Round 2: invertebrate FAIR/ML datasets | 04–05.07.2025 | Google Dataset Search, Zenodo, Dryad, Harvard Datavers | invertebrate AND (tracking OR annotation OR “open dataset”) AND (behavior OR video) | 598 | 166 | Queries adapted per platform syntax. |
| Round 3: model organism generic datasets | 04–06.07.2025 | Google Dataset Search, Zenodo, Dryad, Harvard Dataverse, IEEE DataPort | “model organism” AND (video dataset OR tracking dataset OR annotation dataset) | 395 | 63 | N/A |
| Round 4: organism-specific deep dive | 04–07.07.2025 | Google Dataset Search, Zenodo, Dryad, Harvard Dataverse, OSF | (Drosophila OR “C. elegans” OR “Galleria mellonella” OR zebrafish) AND (movement OR tracking OR behavior) | 700+ | 200+ | Used to reduce false negatives; exact counts reported in Supplementary Data File S6. |
