## Supplemental Data 4 for "Systematic Survey of Public Datasets for Behavioral Research in Invertebrate Models: Toward FAIR and Standardized Data Sharing"

**Supplementary Table S8**. Definitions and ordinal scoring thresholds (1–3) for Stage 2 technical quality dimensions.

| Technical dimension | Rating 1 (low) – threshold | Rating 2 (average) – threshold | Rating 3 (high) – threshold |
|---|---|---|---|
| **Usability** | Data difficult to use without major reconstruction/repair. Deficiencies in structure or files hinder analysis. Unfriendly format (e.g., proprietary without parsers). No clear directory organization. | Data usable after moderate preprocessing. Directory structure partially clear. Minor gaps or inconsistencies exist but do not block analysis. Format conversion required but feasible. | Data ready for use out of the box. Consistent, logical file and directory structure. Open, standard formats. Minimal preprocessing. Examples of loading/analysis included. |
| **Annotation richness** | No behavioral/position/ROI annotations or only very poor ones (e.g., single aggregate metrics). No behavior class labels; difficult to train supervised models. | Partial annotations (e.g., tracking without behavior classes, labels only for part of the data, limited number of classes, unclear label mapping). Enables ML after additional processing. | Rich, well-described behavioral annotations for most data. Clear label definitions and complete annotation files. Ready for training/benchmarking. |
| **Technical quality** | Numerous problems: inconsistent names, missing/damaged files, archive errors, mismatches between metadata and data. Poor recording/tracking quality or missing acquisition parameters. | Overall acceptable quality with some minor technical issues (isolated gaps, occasional tracking errors). Acquisition parameters partially reported; requires user-side verification. | High data quality without significant technical problems. Stable files; author-side quality control evident. Acquisition and processing parameters clearly described. |
| **AI-readiness** | Data unsuitable for AI: lack of raw data and/or training-enabling annotations; formats difficult to process automatically; no versioning/provenance. | Partially AI-ready: raw data and/or annotations available but require substantial preparation (cleaning, harmonization, label acquisition). No metadata standard or only partial code support. | Data prepared specifically for AI: raw data plus training annotations, open formats, stable versions, readable metadata. Attached code/pipeline or usage examples. Easy to reuse in benchmarks. |
