## Supplementary Table S9. Stage 2 ordinal scores and inter-rater agreement per dataset. (1) for "Systematic Survey of Public Datasets for Behavioral Research in Invertebrate Models: Toward FAIR and Standardized Data Sharing"

Two reviewers scored four technical dimensions on a 1–3 ordinal scale (R1, R2). Mean scores are averages across dimensions. Cohen's κ was computed per dataset across the four dimensions; negative κ indicates agreement below chance.

| No. | Dataset name (full) | Usability R1 | Usability R2 | Annotation richness R1 | Annotation richness R2 | Technical quality R1 | Technical quality R2 | AI-readiness R1 | AI-readiness R2 | Mean score R1 | Mean score R2 | Mean overall | Cohen's κ | κ category (Landis–Koch) | Technical issues / notes |
|---|---|---|---|---|---|---|---|---|---|---|---|---|---|---|---|
| 1 | WormSwin: C. elegans Video Datasets | 3 | 2 | 1 | 2 | 3 | 3 | 3 | 2 | 2.50 | 2.25 | 2.38 | 0.08 | Slight (0.00–0.20) | None observed |
| 2 | Statistical analysis and dataset for: Three-dimensional body reconstruction enables quantification of liquid consumption in small invertebrates | 2 | 3 | 2 | 2 | 2 | 2 | 3 | 2 | 2.25 | 2.25 | 2.25 | −0.33 | Poor (<0) | NumPy 2.x compatibility error (modules compiled under NumPy 1.x). |
| 3 | Concatenated planarian behavioral barcodes | 1 | 2 | 1 | 3 | 3 | 3 | 1 | 2 | 1.50 | 2.50 | 2.00 | 0.14 | Slight (0.00–0.20) | None observed |
| 4 | Differential neuroanatomical, neurochemical, and behavioral impacts of early-age isolation in a eusocial insect | 1 | 1 | 2 | 2 | 3 | 3 | 2 | 2 | 2.00 | 2.00 | 2.00 | 1.00 | Almost perfect (>0.80) | None observed |
| 5 | Mantis shrimp locomotion: coordination and variation of hybrid metachronal swimming | 3 | 3 | 3 | 3 | 3 | 3 | 2 | 3 | 2.75 | 3.00 | 2.88 | 0.00 | Slight (0.00–0.20) | None observed |
| 6 | Data from: Multifaceted and extensive behavioral trajectories of genomically diverse Drosophila lines | 2 | 2 | 2 | 3 | 2 | 2 | 3 | 3 | 2.25 | 2.50 | 2.38 | 0.50 | Moderate (0.41–0.60) | No ready behavioral labels; possible tracking–video desynchronization. |
| 7 | Miniature linear and split-belt treadmills reveal mechanisms of adaptive motor control in walking Drosophila | 2 | 3 | 2 | 2 | 3 | 3 | 2 | 2 | 2.25 | 2.50 | 2.38 | 0.50 | Moderate (0.41–0.60) | None observed |
| 8 | Asynchronous haltere input drives specific wing and head movements in Drosophila | 2 | 2 | 2 | 2 | 3 | 2 | 3 | 3 | 2.50 | 2.25 | 2.38 | 0.50 | Moderate (0.41–0.60) | Full video set (~84 GB) could not be downloaded due to storage limits. |
| 9 | 2D Tracking and 3D reconstruction of legs from tethered walking Drosophila | 2 | 2 | 2 | 2 | 3 | 2 | 2 | 2 | 2.25 | 2.00 | 2.12 | 0.00 | Slight (0.00–0.20) | None observed |
| 10 | Crawling Trajectories of Drosophila Larvae Responding to Vibration | 3 | 3 | 3 | 3 | 1 | 2 | 2 | 2 | 2.25 | 2.50 | 2.38 | 0.56 | Moderate (0.41–0.60) | Requires MAGAT Analyzer to open .bin files; installation issues; limited documentation. |
| 11 | Individual flight headings for Drosophila melanogaster in response to different wavelengths of light | 1 | 2 | 1 | 2 | 3 | 1 | 1 | 1 | 1.50 | 1.50 | 1.50 | 0.20 | Slight (0.00–0.20) | None observed |
| 12 | Neural mechanisms to incorporate visual counterevidence in self movement estimation | 2 | 2 | 1 | 1 | 3 | 1 | 2 | 2 | 2.00 | 1.50 | 1.75 | 0.60 | Moderate (0.41–0.60) | None observed |
| 13 | Zebrafish Seizure Behavior Classification Dataset and Trained ML Models | 3 | 3 | 2 | 2 | 2 | 2 | 3 | 3 | 2.50 | 2.50 | 2.50 | 1.00 | Almost perfect (>0.80) | Valid annotations only for baseline group; other conditions have misaligned points. |
| 14 | Chloride-dependent mechanisms of multimodal sensory discrimination and nociceptive sensitization in Drosophila | 2 | 3 | 3 | 3 | 2 | 2 | 2 | 2 | 2.25 | 2.50 | 2.38 | 0.50 | Moderate (0.41–0.60) | None observed |
| 15 | Dimensionality of locomotor behaviors in developing C. elegans | 2 | 2 | 2 | 2 | 3 | 1 | 2 | 2 | 2.25 | 1.75 | 2.00 | 0.43 | Moderate (0.41–0.60) | None observed |
| 16 | Datasets and models for "An improved neural network model enables worm tracking in challenging conditions and increases signal-to-noise ratio in phenotypic screens" | 3 | 3 | 3 | 3 | 2 | 2 | 2 | 3 | 2.50 | 2.75 | 2.62 | 0.50 | Moderate (0.41–0.60) | Very large ZIP archive; download/unpacking issues. |
| 17 | Locomotion of naive Drosophila larvae depending on age | 1 | 1 | 2 | 2 | 2 | 2 | 1 | 1 | 1.50 | 1.50 | 1.50 | 1.00 | Almost perfect (>0.80) | Tracking overlay on video shows frequent tracking errors. |
| 18 | Zebrafish larvae exploration and aversive chemotaxis dataset | 2 | 2 | 2 | 2 | 3 | 3 | 2 | 3 | 2.25 | 2.50 | 2.38 | 0.50 | Moderate (0.41–0.60) | None observed |
| 19 | Sensory neurons couple arousal and foraging decisions in C. elegans | 2 | 2 | 2 | 2 | 3 | 3 | 2 | 2 | 2.25 | 2.25 | 2.25 | 1.00 | Almost perfect (>0.80) | None observed |
| 20 | EigenWorms Dataset | 2 | 2 | 2 | 2 | 2 | 2 | 2 | 3 | 2.00 | 2.25 | 2.12 | 0.00 | Slight (0.00–0.20) | .arff file contains unsupported attribute type; conversion to NUMERIC required. |
| 21 | Scalable Apparatus to Measure Posture and Locomotion (SAMPL) | 1 | 1 | 1 | 1 | 1 | 1 | 1 | 2 | 1.00 | 1.25 | 1.12 | 0.00 | Slight (0.00–0.20) | OSF download extremely slow / did not complete. |
| 22 | Automated multimodal imaging of Caenorhabditis elegans behavior | 1 | 1 | 1 | 2 | 1 | 1 | 1 | 1 | 1.00 | 1.25 | 1.12 | 0.00 | Slight (0.00–0.20) | Data link (Dropbox) inactive; only code repository available. |
| 23 | A reductionist paradigm for high-throughput behavioural fingerprinting in Drosophila | 1 | 1 | 1 | 1 | 1 | 1 | 1 | 2 | 1.00 | 1.25 | 1.12 | 0.00 | Slight (0.00–0.20) | Complex installation (Matlab+R), low code quality; pipeline could not be run. |
