## Supplemental Data 5 for "Systematic Survey of Public Datasets for Behavioral Research in Invertebrate Models: Toward FAIR and Standardized Data Sharing"

**Supplementary Table S10.** Data extraction template and coding scheme for dataset evaluation (RQ1–RQ4).

| Domain | Variable (pole w arkuszu) | Description / what is recorded | Coding / allowed values |
|---|---|---|---|
| **Identification** | Dataset ID | Unique working identifier of the dataset | DS01…DSn (working IDs; independent of numbers in result tables) |
| | Dataset title | Name of the dataset | Text |
| | DOI / persistent link | Persistent identifier or stable link to the dataset repository | DOI / URL / handle / accession |
| | Repository / platform | Hosting repository/platform and its general classification | Enum: Zenodo, Dryad, Figshare, Dataverse, OSF, PANGAEA, GitHub-only, institutional server, other (specify); plus type: general-purpose / specialized |
| | Version retrieved & retrieval date | Latest stable version downloaded for analysis and date of download | Version string (e.g., v1/v2/tag/commit) + YYYY-MM-D |
| **Organism scope (RQ1/2)** | Model organism | Model species (full name) | Text, controlled dictionary |
| | Organism category | Taxonomic category used in synthesis | Enum: C. elegans / Drosophila / Planaria / Galleria / Zebrafish / Other invertebrates |
| | Life stage / strain / sex | Developmental stage, strain, sex (if reported) | Text / NR |
| **Data modality & structure (RQ1)** | Data modality | Which data types are present in the deposit | Multi-select: video, images, tracking/trajectories, time-series, electrophysiology/neural, chemical, 3D recon, ML models, workflows, other |
| | Raw vs derived data | Whether raw recordings and/or processed derivatives are present | Enum: raw only / derived only / both / NR |
| | File formats | All file formats present in the repository | Multi-select: mp4/avi/bin, tiff/png/jpg, csv/tsv/xlsx, hdf5/nwb, mat, json/yaml/xml, arff, other |
| | Folder organization | Directory/file organization pattern | Enum: flat / hierarchical-by-subject / by-experiment / standard-like (e.g., NWB/BIDS) / ad hoc / NR |
| | Internal consistency notes | Evidence of missing files, inconsistent naming, corrupted archives, etc. | Yes / No + short note |
| **Behavior annotations (RQ1)** | Behavioral annotations present | Presence of behavior/pose/ROI annotations | Yes / No / Partial (subset only or pose-only without behavior classes) |
| | Number of behavior classes | Number of behavioral labels/classes | Number / Continuous / NR |
| | Annotation method | How annotations/labels were generated | Enum: manual / semi-automatic / automatic / crowdsourced / NR |
| | Domain of behaviors | Behavioral domains covered | Multi-select: locomotion, foraging, defensive, social, reproductive, sleep/quiescence, other |
| **Acquisition resolution (RQ1/2)** | Temporal resolution | Sampling rate / fps / Hz; session length if specified | Number + unit / NR |
| | Spatial resolution | px/mm, μm/px, 2D/3D tracking level if reported | Text/number + unit / NR |
| **Scale** | Number of individuals | Number of individuals represented | Number / Range / NR |
| | Number of experiments/sessions | Number of sessions/experiments | Number / Approx / NR |
| **Metadata (RQ2)** | Technical documentation present | README/protocol/codebook/pipeline availability | Yes / Partial (incomplete or paper-only) / No |
| | Machine-readable metadata | Whether metadata are machine-readable (not only | Yes (specify format: json/yaml/xml/tsv/…) / No (human-readable only) / NR |
| | Metadata standard / ontology | Use of formal standards/ontologies | Enum: none / custom / partial standard (specify) / named standard (specify, e.g., NWB, NBO, ZECO) / NR |
| **Sharing & FAIR (RQ3/4)** | License type | Dataset license clarity and type | Enum: CC-BY, CC0, MIT, GPL, Apache, CC-BY-NC-*, none/NR, restricted/request-only |
| | Access level | Dataset accessibility category | Enum: open / registration / request-only / paywalled |
| | Persistent identifier present | Presence and type of PID | Yes / No + type (DOI/handle/accession) |
| | FAIR statement/indicators | FAIR-relevant elements visible at dataset/repository level | Checklist: PID, searchable metadata, open formats, license clarity, provenance/versioning, community standard |
| **AI-readiness scoring** | AI-readiness label (pre-score) | Overall qualitative assessment of ML readiness before ordinal scoring | Enum: yes / partial / no |
| | Usability score (1–3) | Ordinal usability rating | 1 / 2 / 3 (thresholds in Supplementary Table S8) |
| | Annotation richness score (1–3) | Ordinal rating of annotation detail | 1 / 2 / 3 (thresholds in Supplementary Table S8) |
| | Technical quality score (1–3) | Ordinal rating of internal technical quality | 1 / 2 / 3 (thresholds in Supplementary Table S8) |
| | AI-readiness score (1–3) | Ordinal overall AI-readiness rating | 1 / 2 / 3 (thresholds in Supplementary Table S8) |
| **Eligibility decision** | Inclusion status | Final Stage 2 eligibility | Enum: included / excluded |
| | Exclusion reason | Brief justification for exclusion if applicable | Controlled categories + free text |
