## Supplementary Table S11. Source-resolved PRISMA 2020 flow summary for Stage 1. for "Systematic Survey of Public Datasets for Behavioral Research in Invertebrate Models: Toward FAIR and Standardized Data Sharing"

| Selection stage | | PubMed | Scopus | Web of Science | IEEE Xplore | ACM Digital Library | Google Scholar | bioRxiv (preprints) | arXiv (preprints) | Reference list screening / citation chasing | Total |
|---|---|---|---|---|---|---|---|---|---|---|---|
| 0 | Total number of records retrieved | 283 | 254 | 805 | 19 | 159 | 200 | 57 | 214 | 67 | 2058 |
| 1 | Number of duplicates | 196 | 210 | 23 | 14 | 22 | 54 | 49 | 187 | 45 | 800 |
| | Total number of records after duplicate removal | 87 | 44 | 782 | 5 | 137 | 146 | 8 | 27 | 22 | **1201** |
| 2 | Number of articles excluded at the abstract-screening stage | 54 | 22 | 750 | 3 | 128 | 140 | 4 | 21 | 18 | 1097 |
| | Total number of articles meeting inclusion criteria (based on abstracts) | 33 | 22 | 32 | 2 | 9 | 6 | 4 | 6 | 4 | **104** |
| 3 | Number of articles excluded at the full-text screening stage | 30 | 19 | 29 | 1 | 8 | 5 | 4 | 6 | 4 | 92 |
| | Total number of articles meeting inclusion criteria (based on full texts) | 3 | 3 | 3 | 1 | 1 | 1 | 0 | 0 | 0 | **12** |
